# Distinct contributions and sensorimotor encoding of PV and FoxP2 neurons in the external globus pallidus during perceptual decision-making

**DOI:** 10.1101/2025.09.03.673900

**Authors:** Xiaotian Pu, Jingyu Ren, Taorong Xie, Kexin Zou, Jingze Xu, Yaping Li, Dechen Liu, Haishan Yao

**Affiliations:** Institute of Neuroscience, State Key Laboratory of Brain Cognition and Brain-inspired Intelligence Technology, Center for Excellence in Brain Science and Intelligence Technology, Chinese Academy of Sciences, Shanghai 200031, China; University of Chinese Academy of Sciences, Beijing 100049, China

## Abstract

The external globus pallidus (GPe), a central nucleus of the basal ganglia (BG), comprises diverse cell types, including PV and FoxP2 neurons. While their roles in movement regulation and sensory processing have been studied, their contributions to perceptual decision-making remain unclear. Using a Go/No-Go visual task in mice, we found that optogenetic activation of either population impaired behavioral performance, whereas only inhibition of PV neurons, but not FoxP2 neurons, disrupted behavior. Electrophysiological recordings revealed that PV and FoxP2 neurons exhibited distinct encoding of visual stimuli, licking movement, and trial outcomes, with a larger fraction of PV neurons positively encoding the reward-predicting (Go) stimulus and the population overall preferring the rewarded over unrewarded outcome. FoxP2 neuron inhibition had minimal effects on spontaneous firing of neurons in the substantia nigra pars reticulata (SNr), a major BG output. By contrast, PV neuron inhibition altered both spontaneous and task-related activity of SNr neurons, reducing their preference for reward-predicting stimulus or unrewarded outcome in a subpopulation-specific manner. These findings indicate that PV and FoxP2 neurons in the GPe differentially influence sensorimotor processing and SNr activity, with PV neurons playing a key role in supporting accurate performance during perceptual decision-making.

## Introduction

The external globus pallidus (GPe), a central nucleus in the basal ganglia (BG), plays a crucial role in motor control and cognitive functions (Courtney and Chan, 2023; Courtney et al., 2023; Dong et al., 2021; Giossi et al., 2024; Gittis et al., 2014; Hegeman et al., 2016; Nambu and Chiken, 2024). Traditionally, the GPe was viewed primarily as a relay nucleus in the indirect pathway of the BG circuit, where its GABAergic neurons receive inhibitory inputs from striatal medium spiny neurons expressing the D2-type dopamine receptors (D2-MSNs) and excitatory inputs from the subthalamic nucleus (STN), and project to STN and the BG output nuclei, the substantia nigra pars reticulata (SNr) and internal globus pallidus (GPi) (Gerfen et al., 1990; Kita, 2007). Emerging evidence, however, has revealed that the GPe is more than a simple relay; it forms extensive connections with all major BG components, as well as with the thalamus and cortex (Courtney et al., 2023; Dong et al., 2021; Fang and Creed, 2024; Giossi et al., 2024; Mastro et al., 2014). Moreover, the GPe consists of diverse cell types that differ in their projection patterns, molecular profiles, electrophysiological characteristics, and functional properties (Courtney et al., 2023; Fang and Creed, 2024; Giossi et al., 2024; Guilhemsang and Mallet, 2024).

Early research identified a dichotomous organization within the GPe: prototypic neurons are tonically active and innervate canonical downstream nuclei such as the STN, and arkypallidal neurons are characterized by lower spike rates and project back to the striatum (Abdi et al., 2015; Dodson et al., 2015; Mallet et al., 2012). Molecularly, GPe neurons are divided into distinct populations, with approximately 50% expressing parvalbumin (PV) and around 20% expressing forkhead box protein P2 (FoxP2) (Cui et al., 2021b; Dodson et al., 2015; Hernandez et al., 2015; Saunders et al., 2016). GPe PV neurons display prototypic properties, innervating the STN, SNr, and parafascicular nucleus (PF) of the thalamus (Hernandez et al., 2015; Mastro et al., 2014). These neurons also contain subpopulations that co-express other molecular markers (Abecassis et al., 2020). In contrast, GPe FoxP2 neurons exhibit arkypallidal properties, exclusively innervating the striatum (Abdi et al., 2015). The FoxP2 neurons are a major subclass of neurons expressing neuronal PAS-domain-protein-1 (Npas1), which make up about 30% of GPe population and do not overlap with PV-expressing neurons (Abecassis et al., 2020; Dodson et al., 2015; Hernandez et al., 2015; Pamukcu et al., 2020). This cellular diversity necessitates a cell-type-specific approach to investigating GPe function.

Behavioral significance of specific GPe cell types has been explored in various studies. In vivo recordings have shown distinct activity patterns of prototypic and arkypallidal GPe neurons during movements (Dodson et al., 2015; Mallet et al., 2016). In healthy mice, optogenetic activation of GPe PV neurons enhances locomotor activity, whereas their inhibition suppresses it (Cui et al., 2021b; Lilascharoen et al., 2021; Pamukcu et al., 2020). Conversely, activation of GPe Npas1 or FoxP2 neurons reduces locomotor speed (Aristieta et al., 2021; Cui et al., 2021b; Glajch et al., 2016; Labouesse et al., 2023; Pamukcu et al., 2020), while Npas1 neuron inhibition increases locomotion (Pamukcu et al., 2020). Combinatorial movement metrics reveal that optogenetic stimulation of GPe FoxP2 neurons induces behavioral changes similar to those of Npas1 activation but opposite to those elicited by PV stimulation (Cui et al., 2021b). Recent studies further highlight the contribution of specific GPe cell types to movement and volitional control in normal mice, as well as to hypokinetic symptoms or dyskinetic behaviors in mouse models of Parkinson’s disease (PD) (Lilascharoen et al., 2021; Luo et al., 2025; Mastro et al., 2017; Pamukcu et al., 2020; Shen et al., 2024; Spix et al., 2021; Xu et al., 2025). Beyond motor control, GPe neurons have also been implicated in non-motor functions (Courtney and Chan, 2023; Dong et al., 2021; Gittis et al., 2014; Schechtman et al., 2016). For example, they encode information related to value, salience, and valence (Arkadir et al., 2004; Farries et al., 2023; Katabi et al., 2023; Kim et al., 2017). GPe arkypallidal neurons participate in movement suppression in a stop-signal task (Mallet et al., 2016) and regulate habitual seeking behaviors (Baker et al., 2023), while GPe PV neurons mediate behavioral punishment (Isett et al., 2023) and control cocaine sensitivity (Beier et al., 2017; Tian et al., 2024), highlighting their important contribution to decision-making processes and related cognitive functions. In anesthetized mice, prototypic and arkypallidal GPe neurons exhibit distinct response patterns to tactile sensory stimulation (Ketzef and Silberberg, 2021), which are attributed to their different input sources from the striatum and STN (Johansson and Ketzef, 2023). Collectively, these studies underscore the integrative role of the GPe in processing diverse motor and non-motor signals. However, it remains unclear how different types of GPe neurons contribute to sensorimotor function and encode task-related signals during perceptual decision-making, a process in which animals use sensory cues to guide appropriate motor responses. Furthermore, studies in PD mouse models have found that manipulating specific cell types in the GPe alters pathological firing patterns in SNr neurons and rescues motor behavior (Mastro et al., 2017; Spix et al., 2021). These findings raise the question of whether, and how, defined GPe cell types modulate spontaneous and task-related activity in SNr neurons under normal conditions. In this study, we investigated the functional roles of GPe PV and FoxP2 neurons during a Go/No-Go visual discrimination task in mice. We found that activation of PV or FoxP2 neurons altered behavior, and inhibition of PV neurons also decreased performance. Optogenetic identification revealed that PV and FoxP2 neurons exhibited distinct patterns of task-related activity. Furthermore, inhibiting PV but not FoxP2 neurons changed spontaneous SNr firing, and PV inhibition differentially modulated stimulus- and outcome-related activity across two subpopulations of putative GABAergic SNr neurons during task engagement. These findings provide insight into the differential roles of GPe PV and FoxP2 neurons in regulating sensorimotor processing during perceptual decision-making.

## Results

### Activation of either PV or FoxP2 neurons impairs Go/No-Go task performance

Head-fixed mice were trained to perform a Go/No-Go visual discrimination task (Liu et al., 2023) (Figure 1A), in which the Go and No-Go stimuli were vertical and horizontal gratings, respectively. The duration of stimulus presentation consisted of a waiting period and an answer period. During the answer period, licking in response to the Go stimulus was rewarded with water (Hit), while licking to the No-Go stimulus was false alarm (FA) and was not rewarded.

**Figure 1.**
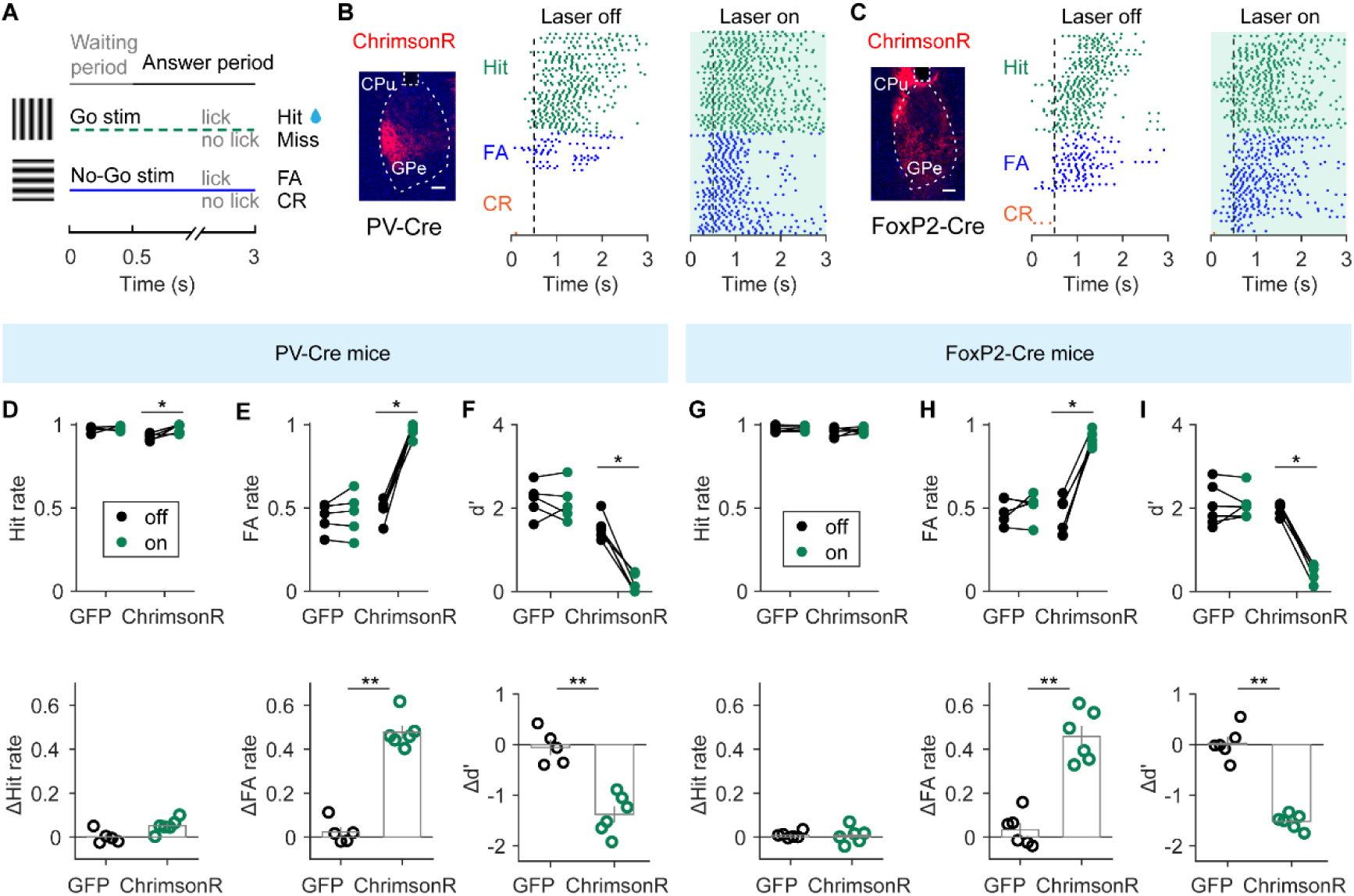
Activation of PV or FoxP2 GPe neurons impairs performance in the Go/No-Go visual task. (A) Schematic of the trial structure. (B) Left, representative fluorescence image showing AAV-FLEX-ChrimsonR expression in the GPe of a PV-Cre mouse. Right, lick rasters from an example PV-Cre mouse during laser-off and laser-on trials. For display purposes, 100 laser-off trials and 100 laser-on trials were shown in the raster plot. Shading, duration of laser stimulation. (C) Left, representative fluorescence image showing AAV-FLEX-ChrimsonR expression in the GPe of a FoxP2-Cre mouse. Right, lick rasters from an example FoxP2-Cre mouse during laser-off and laser-on trials (100 trials were shown for each condition). Shading, duration of laser stimulation. (D) Top, Hit rates in laser-off and laser-on trials for control (n = 5) and ChrimsonR (n = 6) PV-Cre mice. * p = 0.031, Wilcoxon signed-rank test. Bottom, comparison of ΔHit rate between control and ChrimsonR PV-Cre mice. p = 0.056, Wilcoxon rank-sum test. (E) Same as (D), but for FA rates in GFP and ChrimsonR PV-Cre mice. Top, * p = 0.031, Wilcoxon signed-rank test. Bottom, ** p = 0.0043, Wilcoxon rank-sum test. (F) Same as (D), but for d′ in GFP and ChrimsonR PV-Cre mice. Top, * p = 0.031, Wilcoxon signed-rank test. Bottom, ** p = 0.0043, Wilcoxon rank-sum test. (G) Top, Hit rates in laser-off and laser-on trials for control (n = 6) and ChrimsonR (n = 6) FoxP2-Cre mice. Bottom, comparison of ΔHit rate between control and ChrimsonR FoxP2-Cre mice. P = 0.97, Wilcoxon rank-sum test. (H) Same as (G), but for FA rates in GFP and ChrimsonR FoxP2-Cre mice. Top, * p = 0.031, Wilcoxon signed-rank test. Bottom, ** p = 0.0022, Wilcoxon rank-sum test. (I) Same as (G), but for d′ in GFP and ChrimsonR FoxP2-Cre mice. Top, * p = 0.031, Wilcoxon signed-rank test. Bottom, ** p = 0.0022, Wilcoxon rank-sum test. Scale bars, 200 μm. Data represent mean ± SEM.

To examine the effect of activating GPe PV or FoxP2 neurons, we used PV-Cre or FoxP2-Cre mice (Figure 1B and C), in which AAV-FLEX-ChrimsonR was unilaterally injected into the anterior GPe. In vivo recordings verified that optogenetic activation of ChrimsonR could increase the firing rates of GPe neurons (Figure 3). We trained the mice to perform the task using visual stimuli presented contralateral to the virus injection site. After stable task performance was achieved, red laser stimulation (635 nm, 0 – 3 s relative to stimulus onset) was applied to the GPe during task execution in laser-on blocks, which were interleaved with laser-off blocks (Figure 1B and C). We found that activation of GPe PV neuron significantly increased the FA rate (p = 0.031, n = 6, Wilcoxon signed-rank test, Figure 1E, top panel) as well as the Hit rate (p = 0.031, n = 6, Wilcoxon signed-rank test, Figure 1D, top panel), with a significant decrease in discriminability (d’) (p = 0.031, n = 6, Wilcoxon signed-rank test, Figure 1F, top panel). In control PV-Cre mice in which AAV-FLEX-EGFP were injected in the GPe, laser stimulation did not affect behavioral performance (p > 0.5, n = 5, Wilcoxon signed-rank test, Figures 1D-F, top panels). Compared to the GFP mice, the ChrimsonR mice showed a significantly larger increase in FA rate and greater reduction in d′ following laser stimulation (p < 0.01, Wilcoxon rank-sum test, Figure 1E and F, bottom panels). Similar to PV neuron activation, activating FoxP2 neurons in the GPe also increased the FA rate and reduced d′ (p < 0.05, Wilcoxon signed-rank test, Figure 1G-I).

Since activating PV neurons during the stimulus presentation period increased lick rates in both Hit and FA trials (Figure 1−figure supplement 1), we next asked whether the increased FA rate could be attributed to a direct facilitation of licking behavior. To test this possibility, we activated PV or FoxP2 neurons during the inter-trial interval (ITI), defined as the period between the end of the answer period of the preceding trial and the onset of the subsequent stimulus. In contrast to stimulus-period activation, activation of PV or FoxP2 neurons during the ITI did not significantly alter licking rates compared to controls (Figure 1−figure supplement 1). These results suggest that the impaired task performance during stimulus-period activation is unlikely to arise from a direct motor effect on licking behavior.

### Inhibition of PV but not FoxP2 neurons disrupts behavioral performance

We next performed experiments to inhibit PV or FoxP2 neurons during the stimulus presentation period, by injecting AAV-DIO-NpHR into the anterior GPe of PV-Cre or FoxP2-Cre mice. In vivo recordings confirmed that activation of NpHR in GPe PV neurons effectively suppressed GPe neuron firing (Figure 2−figure supplement 1). Similarly, in vitro recordings showed that NpHR activation in GPe FoxP2 neurons reduced spiking activity evoked by current injection (Figure 2−figure supplement 1). Compared to control mice, inhibition of PV neurons significantly increased the FA rate and decreased d′ (p < 0.001, Wilcoxon rank-sum test, Figure 2A-D), whereas inhibition of FoxP2 neurons had no effect on behavioral performance (p > 0.5, Wilcoxon rank-sum test, Figure 2E-H). Lick rate analysis showed that inhibiting PV neurons during the stimulus presentation period increased licking in FA trials (Figure 2−figure supplement 1), whereas inhibiting FoxP2 neurons had no effect on licking (Figure 2−figure supplement 1). To test whether the increased FA rate was due to a direct influence on licking behavior, we inhibited PV neurons during the ITI. Under this condition, the NpHR group did not show an increased lick rate compared to controls (Figure 2−figure supplement 1). These results suggest that the increased FA rate and decreased d′ caused by PV neuron inhibition were not attributable to direct modulation of licking movements.

**Figure 2.**
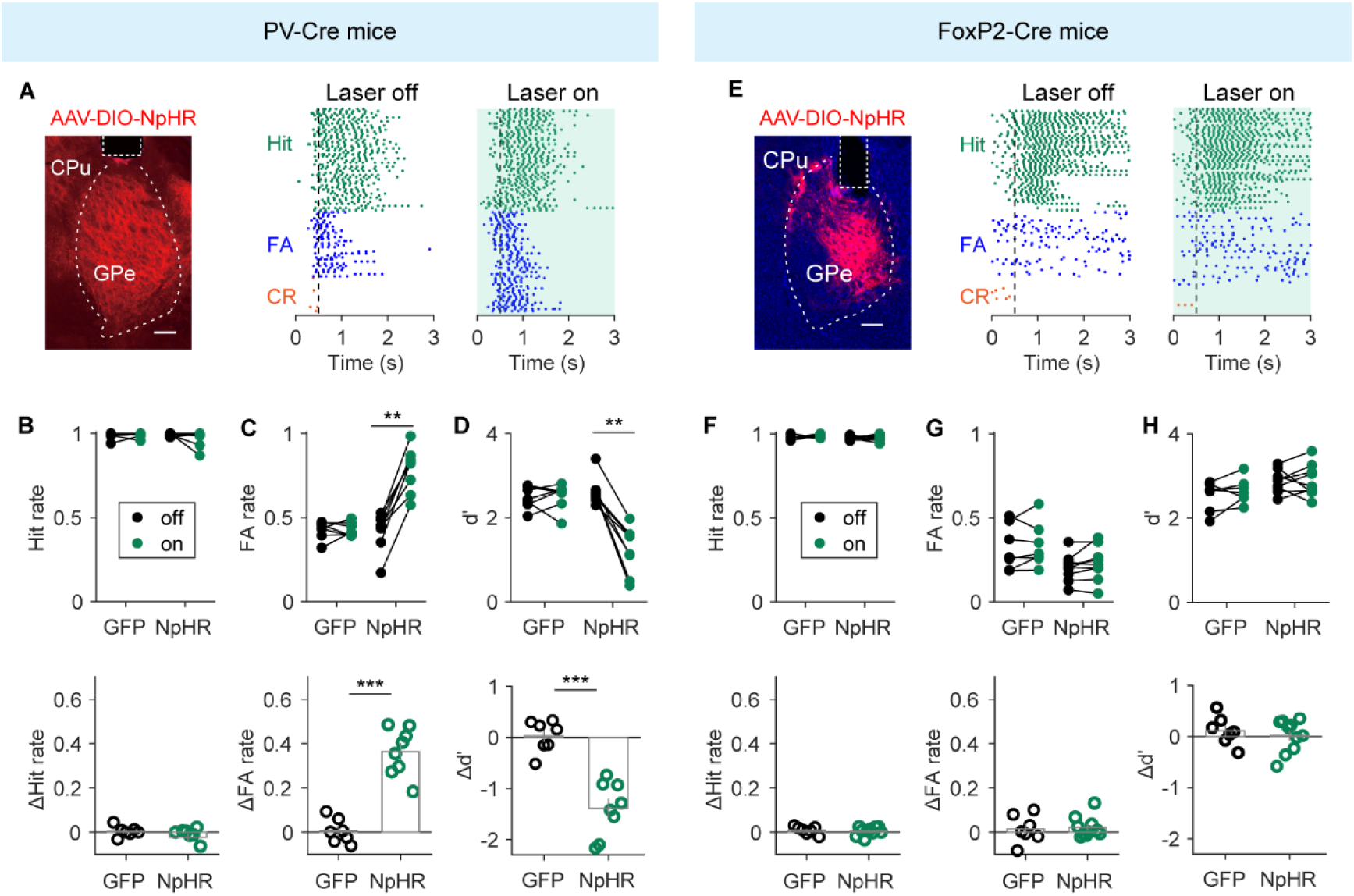
Inactivation of PV but not FoxP2 GPe neurons impairs performance in the Go/No-Go visual task. (A) Left, representative fluorescence image showing AAV-DIO-NpHR expression in the GPe of a PV-Cre mouse. Right, lick rasters from an example PV-Cre mouse in laser-off and laser-on trials. For display purposes, 100 laser-off trials and 100 laser-on trials were shown in the raster plot. Shading, duration of laser stimulation. (B) Top, Hit rates in laser-off and laser-on trials for control (n = 7) and NpHR (n = 8) PV-Cre mice. Bottom, comparison of ΔHit rate between control and NpHR PV-Cre mice. p = 0.34, Wilcoxon rank-sum test. (C) Same as (B), but for FA rates in GFP and NpHR PV-Cre mice. Top, ** p = 0.008, Wilcoxon signed-rank test. Bottom, *** p = 3.11×10^-4^, Wilcoxon rank-sum test. (D) Same as (B), but for d′ in GFP and NpHR PV-Cre mice. Top, ** p = 0.008, Wilcoxon signed-rank test. Bottom, *** p = 3.11×10^-4^, Wilcoxon rank-sum test. (E) Left, representative fluorescence image showing AAV-DIO-NpHR expression in the GPe of a FoxP2-Cre mouse. Right, lick rasters from an example FoxP2-Cre mouse in laser-off and laser-on trials (100 trials were shown for each condition). Shading, duration of laser stimulation. (F) Top, Hit rates in laser-off and laser-on trials for control (n = 7) and NpHR (n = 10) FoxP2-Cre mice. Bottom, p = 0.87, Wilcoxon rank-sum test. (G) Same as (F), but for FA rates in GFP and NpHR FoxP2-Cre mice. Bottom, p = 0.96, Wilcoxon rank-sum test. (H) Same as (F), but for d′ in GFP and NpHR FoxP2-Cre mice. Bottom, p = 0.74, Wilcoxon rank-sum test. Scale bars, 200 μm. Data represent mean ± SEM.

**Figure 3.**
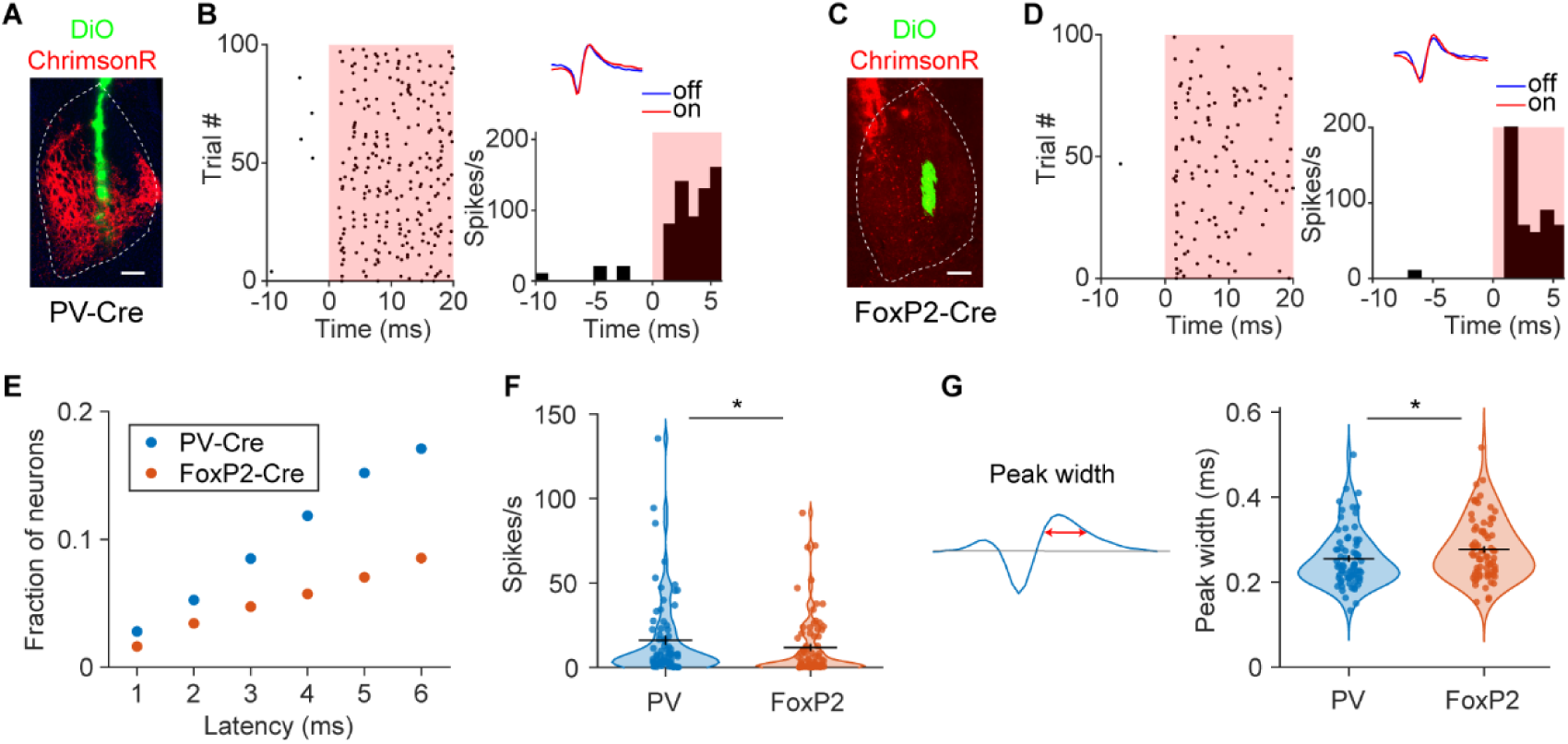
Identification of PV or FoxP2 GPe neurons. (A) Fluorescence image showing AAV-FLEX-ChrimsonR-tdTomato expression in the GPe of a PV-Cre mouse. Green, electrode track labeled with DiO. (B) Spike rasters (left) and PSTH (right) of an identified PV neuron. Time 0 marks laser onset. Red shading, laser stimulation. Inset, average spike waveforms recorded during laser-off and laser-on trials. (C) Fluorescence image showing AAV-FLEX-ChrimsonR-tdTomato expression in the GPe of a FoxP2-Cre mouse. Green, electrode track labeled with DiO. (D) Spike rasters (left) and PSTH (right) of an identified FoxP2 neuron (format as in B). (E) Fraction of identified PV and FoxP2 GPe neurons using latency threshold ranging from 1 to 6 ms. PV neurons were identified from a total of 895 units in 50 sessions of 8 PV-Cre mice. FoxP2 neurons were identified from a total of 1606 units in 85 sessions of 14 FoxP2-Cre mice. (F) Comparison of spike rates between PV neurons (n = 76) and FoxP2 GPe neurons (n = 76) identified using a 3-ms latency threshold. * p = 0.048, Wilcoxon rank-sum test. (G) Comparison of spike peak width (the width at half-maximum of the waveform peak amplitude) between PV and FoxP2 GPe neurons identified using a 3-ms latency threshold. * p = 0.036, Wilcoxon rank-sum test. Scale bars, 200 μm.

These results indicate that manipulating GPe PV neuron activity during the stimulus presentation period, either through activation or inhibition, disrupts behavioral performance in the Go/No-Go task. In contrast, although activation of FoxP2 neurons increased FA rate, inhibition of FoxP2 neurons did not significantly affect task performance. Thus, our results highlight an important role for GPe PV neurons in supporting accurate sensory-guided behavior.

### Optogenetic tagging of GPe PV and FoxP2 neurons

To identify GPe PV and FoxP2 neurons for subsequent analysis of task-related activity, we performed optogenetic tagging with an optotrode during electrophysiological recordings in the anterior GPe (Figure 3−figure supplement 1) in PV-Cre or FoxP2-Cre mice, in which AAV-FLEX-ChrimsonR had been injected (Figure 3A and C). Neurons were identified as ChrimsonR-expressing based on two criteria (Chen et al., 2021; Lee et al., 2019): (1) significant activation by laser stimulation with a short latency (within 6 ms), and (2) a Pearson’s correlation coefficient > 0.95 between laser-evoked and spontaneous spike waveforms (Figure 3B and D).

Depending on the latency threshold applied (1−6 ms), we identified 25−153 PV neurons out of 895 recorded units in PV-Cre mice, and 26−137 FoxP2 neurons out of 1606 units in FoxP2-Cre mice (Figure 3E and Figure 3−figure supplement 1). In control PV-Cre and FoxP2-Cre mice without virus injection, no neurons satisfied the criteria for optogenetic identification at any latency threshold between 1 and 6 ms (Figure 3–figure supplement 1). At thresholds between 2 and 4 ms, identified PV neurons exhibited higher spontaneous firing rates and narrower spike waveform peak widths compared to FoxP2 neurons (Figure 3F and G, and Figure 3−figure supplement 1), consistent with previous reports (Aristieta et al., 2021; Dodson et al., 2015; Koene et al., 2025). For subsequent analyses, we adopted a 3 ms latency threshold, which provided a more stringent classification of ChrimsonR-expressing cells while retaining a sufficient number of neurons for statistical analysis.

### Distinct task-related activity patterns in GPe PV and FoxP2 neurons

We next analyzed GPe PV and FoxP2 neurons that were task-modulated, defined as neurons whose firing rates were significantly modulated by at least one main effect or interaction of stimulus, action, or task epoch in a three-way ANOVA (p < 0.01; see Materials and methods). Example spike rasters and the corresponding peristimulus time histograms (PSTHs) from representative neurons are shown in the top and middle rows of Figure 4B and C, respectively. Across the population, these task-modulated neurons exhibited diverse firing patterns that were time-locked to key task events, including stimulus onset, lick-bout onset and offset, and outcome onset. To disentangle the contributions of each task event to neuronal activity, we fitted the responses of task-modulated neurons using a temporal-kernel generalized linear model (GLM) (Pinto and Dan, 2015). The model included regressors for sensory stimuli (Go and No-Go), licking actions (lick-bout onset and offset), and outcomes (Hit, CR, and FA) (Figure 4A), each represented by a set of time-lagged predictors. This approach enabled the GLM coefficients to capture the temporal dynamics of task-related neuronal responses (see Materials and methods). For the example neurons (Figure 4B and C), the temporal profiles of GLM coefficients (bottom row) closely matched the corresponding PSTHs (middle row). Model performance was evaluated by computing the Pearson’s correlation coefficient (CC) between the predicted and actual Z-scored firing rates on held-out test data. CC values did not differ significantly between PV and FoxP2 neurons (p = 0.66, Wilcoxon rank-sum test; PV: 0.29 ± 0.02, n = 51; FoxP2: 0.30 ± 0.02, mean ± SEM, n = 52) (Figure 3−figure supplement 1), indicating comparable model fits across cell types. Moreover, the behavioral discrimination performance of PV-Cre and FoxP2-Cre mice during these electrophysiological recordings did not differ significantly (Figure 3−figure supplement 1).

**Figure 4.**
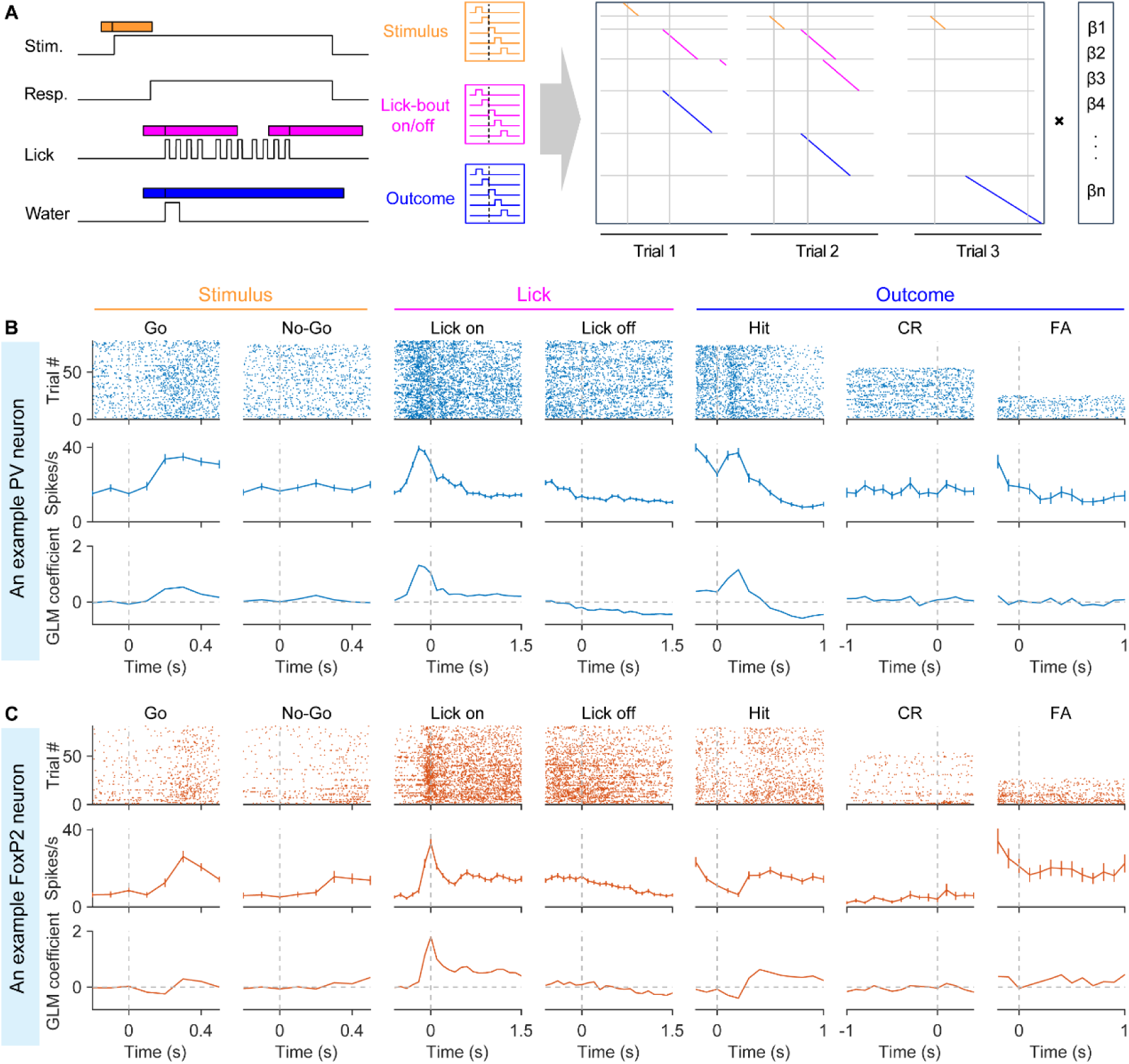
GLM analysis of task-related activity in example GPe neurons. (A) Schematic of the GLM framework, showing task-related variables and their corresponding weights. (B) Spike rasters (top), PSTHs (middle), and GLM coefficients (bottom) from an example PV neuron, aligned to different task events: stimulus onset (Go and No-Go), lick-bout onset and offset, and outcome events (Hit: reward onset, CR: end of response window, FA: onset of sound cue triggered by the first lick in FA trials). For display purposes, 80 Go trials and 80 No-Go trials were shown in the rasters and PSTHs; GLM coefficients were computed from all trials. (C) Same as (B), but for an example FoxP2 neuron.

Averaging the temporal-kernel GLM coefficients across neurons within each cell type revealed significant differences in encoding dynamics between PV and FoxP2 neurons for the Go stimulus, licking events, and outcome events (two-way ANOVA with mixed design followed by Tukey’s multiple comparisons test, p < 0.05 for each event, Figure 5A). To examine these differences at the single-neuron level, we averaged the temporal-kernel GLM coefficients within a defined time window for each event for each neuron, and then compared the mean coefficients between PV and FoxP2 neurons (Figure 5B). PV neurons exhibited significantly higher mean coefficients for the Go stimulus than FoxP2 neurons (Figure 5B, p = 8.25×10^-3^, Wilcoxon rank-sum test), suggesting stronger positive encoding of reward-predicting sensory cue. By contrast, for lick-bout offset, PV neurons had negative mean coefficients that were significantly lower than those of FoxP2 neurons (Figure 5B, p = 0.015, Wilcoxon rank-sum test), indicating stronger negative encoding of lick-bout termination in PV neurons. Across task events, individual neurons displayed both positive and negative coefficients (Figure 5B), reflecting heterogeneity in encoding within each cell type.

**Figure 5.**
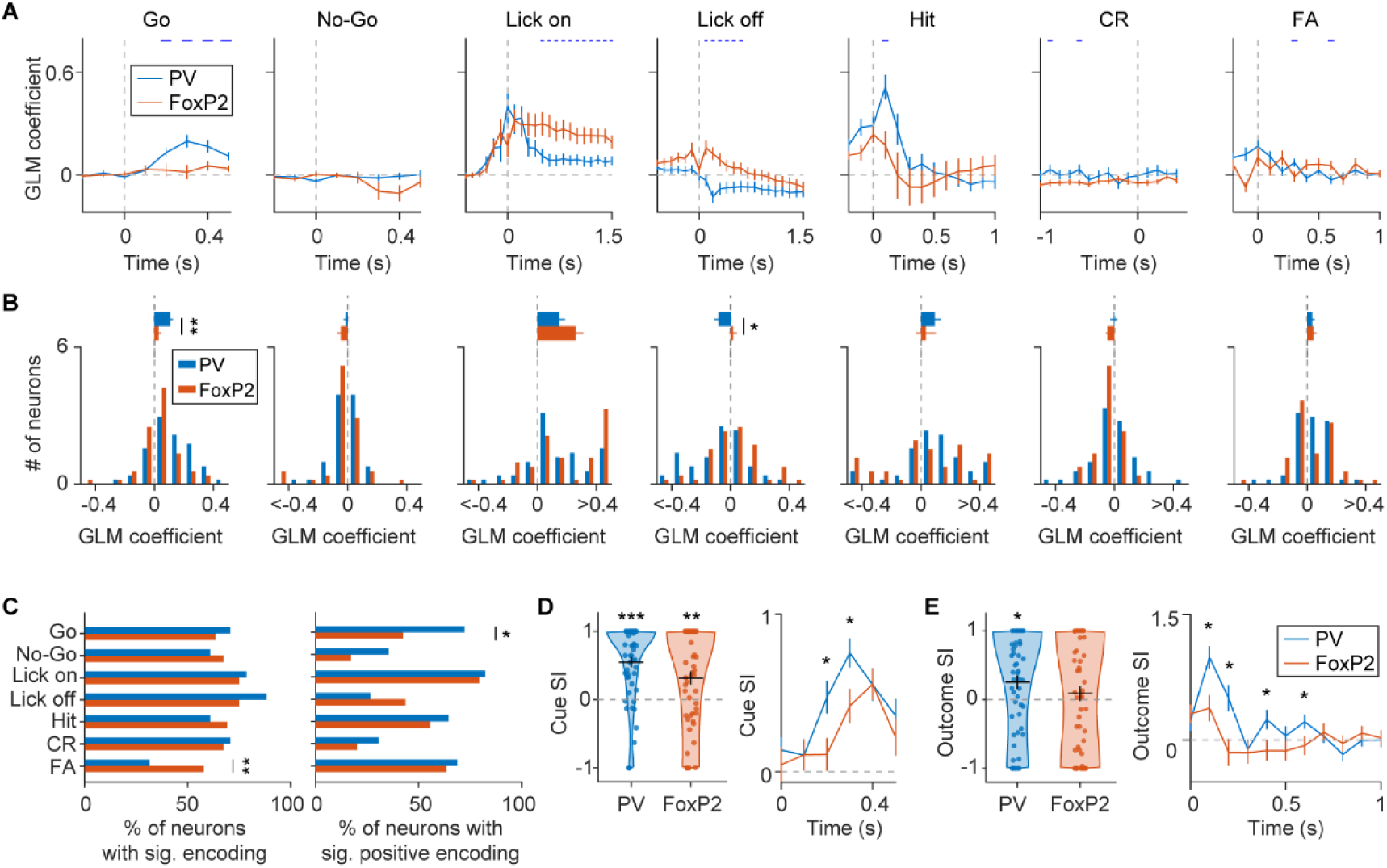
Comparison of GLM coefficients between task-modulated PV and FoxP2 GPe neurons. (A) Temporal profiles of GLM coefficient averaged across task-modulated PV neurons (n = 51) or FoxP2 neurons (n = 52) for each regressor. Horizontal ticks indicate time points where GLM coefficients differed significantly between PV and FoxP2 neurons (p < 0.05), determined by a two-way ANOVA with mixed design followed by Tukey’s multiple comparisons test. For each regressor, the two-way ANOVA was applied only to time points within specified time windows: Go or No-Go stimulus, 0 – 0.5 s; lick-bout onset, 0 – 1.5 s; lick-bout offset, 0 – 1.5 s; Hit, 0 – 1 s; CR, −1 – 0 s (with time zero defined as the end of the answer period); and FA, 0 – 1 s. (B) Distribution of the mean GLM coefficients averaged over the same time windows specified in (A). For Go stimulus, p = 0.004, two-sample Kolmogorov-Smirnov test. For lick-bout offset, p = 0.043, two-sample Kolmogorov-Smirnov test. Top, average of GLM coefficients, * p < 0.05, ** p < 0.01, Wilcoxon rank-sum test. (C) Left, proportion of PV and FoxP2 neurons showing significant encoding for each task event. Right, among neurons with significant encoding, proportion of PV and FoxP2 neurons exhibiting significantly positive (or negative) encoding for each event. * p < 0.05, ** p < 0.01, Fisher’s exact test. (D) Left, cue SI of PV and FoxP2 GPe neurons. ** p < 0.01, *** p < 0.001, Wilcoxon signed-rank test. Right, temporal dynamics of cue SI for PV and FoxP2 neurons. * p < 0.05, two-way ANOVA with mixed design (F_(1, 101)_ = 4.03, p = 0.047) followed by Tukey’s multiple comparisons test. (E) Left, outcome SI of PV and FoxP2 GPe neurons. * p < 0.05, Wilcoxon signed-rank test. Right, temporal dynamics of outcome SI for PV and FoxP2 neurons. * p < 0.05, two-way ANOVA with mixed design (F_(10, 1010)_ = 4.31, p_interaction_ = 0.0018) followed by Tukey’s multiple comparisons test. PV neurons and FoxP2 neurons were identified using a latency threshold of 3 ms. Data represent mean ± SEM.

To quantify the fraction of neurons with significant positive or negative encoding of each task event, we assessed the contribution of each event by comparing full and reduced models lacking that event. An event was considered to be significantly encoded by a neuron if exclusion of that event resulted in a significant reduction in model performance (see Materials and methods). Across the seven task events, the proportions of neurons with significant encoding did not differ between PV and FoxP2 populations for six events (Figure 5C, p > 0.05, Fisher’s exact test). However, for the FA event (unrewarded outcome), FoxP2 neurons exhibited a significantly higher fraction of cells with significant encoding compared to PV neurons (Figure 5C, p = 0.0099, Fisher’s exact test). Conversely, among neurons that significantly encoded the Go stimulus, a higher fraction of PV neurons had positive coefficients compared with FoxP2 neurons (Figure 5C, p = 0.016, Fisher’s exact test), suggesting a stronger bias toward reward-predictive sensory cue in the PV population.

To further characterize stimulus preference at the single-neuron level, we computed a cue selectivity index (cue SI) from temporal-kernel GLM coefficients, defined as (β_Go_ - β_No-Go_)/(|β_Go_| + |β_No-Go_|), in which β_Go_ and β_No-Go_ denote the average coefficients within the 0 – 0.5 s window following stimulus onset in Go and No-Go trials, respectively. Cue SI values ranged from −1 to 1, reflecting both the preferred stimulus identity and the strength of preference. Both PV and FoxP2 neurons exhibited cue SI values significantly greater than 0 (p < 0.01, Wilcoxon signed-rank test, Figure 5D, left), indicating a population-level preference for the Go stimulus. Across the population, cue SI values did not differ significantly between PV and FoxP2 neurons (p = 0.16, Wilcoxon rank-sum test, Figure 5D, left). However, a sliding-window analysis revealed that PV neurons displayed significantly higher cue SI values within the first 0.3 s following stimulus onset (F_(1, 101)_ = 4.03, p = 0.047, two-way ANOVA with mixed design, Figure 5D, right), suggesting a stronger early preference for reward-predicting cue compared with FoxP2 neurons.

Using the same temporal-kernel GLM framework applied for cue SI, we next quantified outcome selectivity at the single-neuron level. We calculated an outcome selectivity index (outcome SI) (Figure 5E), defined as (β_Hit_ - β_FA_)/(|β_Hit_| + |β_FA_|), where β_Hit_ and β_FA_ represent the average coefficients in the 0 – 1 s window following outcome onset in Hit and FA trials, respectively. Outcome SI values ranged from −1 to 1, with positive values reflecting stronger responses to rewarded (Hit) outcome and negative values indicating stronger responses to unrewarded (FA) outcome. PV neurons exhibited significantly positive outcome SI values (p = 0.016, Wilcoxon signed-rank test, Figure 5E, left), whereas the outcome SI of FoxP2 neurons did not differ significantly from zero (p = 0.43, Wilcoxon signed-rank test, Figure 5E, left). A sliding-window analysis over the 1 s following outcome onset further revealed that PV neurons displayed significantly higher outcome SI values than FoxP2 neurons at multiple time points during the early outcome period (F_(10, 1010)_ = 4.31, p_interaction_ = 0.0018, two-way ANOVA with mixed design, Figure 5E, right), indicating a stronger preference for rewarded over unrewarded outcomes in PV neurons.

To validate the GLM results, we plotted the Z-scored PSTHs averaged across all PV neurons and across all FoxP2 neurons, aligned to each task event (Figure 5-figure supplement 1). The average activity profiles recapitulated the differences observed in the temporal-kernel GLM coefficients between the two cell types. Similar differences in the fractions of neurons with significant encoding of task events, cue SI, and outcome SI between PV and FoxP2 neurons were also observed when neurons were identified using either a 2 ms latency threshold (Figure 5-figure supplement 2) or a 4 ms threshold (Figure 5-figure supplement 3), indicating that these effects were robust across classification criteria.

Together, these analyses demonstrate that both PV and FoxP2 neurons encoded task-relevant events, but exhibited distinct population- and single-neuron-level patterns. Across the entire population, FoxP2 neurons contained a higher fraction of cells that significantly encoded unrewarded outcome. Among neurons with significant encoding, PV neurons preferentially represented reward-predictive Go stimulus and rewarded outcome.

### Impact of GPe PV neuron activity on task-related responses in the SNr

Previous studies have shown that manipulating GPe PV neurons can modulate pathological burst spiking in SNr neurons in a mouse model of PD (Mastro et al., 2017). However, how distinct GPe neuronal subtypes influence SNr activity under physiological conditions remains unclear.

We first examined the effect of inhibiting GPe PV or FoxP2 neurons on the spontaneous firing of SNr neurons. We expressed NpHR in either PV or FoxP2 neurons in the GPe and performed extracellular recordings in the medial and central SNr (Figure 6−figure supplement 1). Putative GABAergic SNr units were identified based on their spike waveforms (Barter et al., 2015) (Figure 6−figure supplement 1), and neurons with baseline firing rates exceeding 5 Hz were included in the analyses. Inactivation of GPe FoxP2 neurons did not significantly alter the spontaneous firing in the majority of putative GABAergic SNr neurons (94.7%, 90/95), with only a few showing changes (Figure 6−figure supplement 1). In contrast, inactivation of GPe PV neurons produced robust effects, with 28.9% (35/121) of SNr neurons exhibiting a significant increase in spontaneous firing rate and another 28.9% (35/121) showing a significant decrease (Figure 6−figure supplement 1).

Given the pronounced effect of GPe PV neuron inhibition on SNr spontaneous firing, we next examined how GPe PV neuron activity influences task-related signals in the SNr. To this end, we combined optogenetic inactivation of GPe PV neurons with extracellular recordings in the SNr during task performance (Figure 6A). In PV-Cre mice, this manipulation significantly increased the FA rate (p = 2.4×10^-4^, Wilcoxon signed-rank test) and decreased d’ (p = 2.4×10^-4^, n = 13 sessions from 4 mice, Wilcoxon signed-rank test, Figure 6B). At the neuronal level, 39.4% (74/188) of putative GABAergic SNr neurons exhibited a significant increase in mean firing rate following GPe PV inactivation, while 46.8% (88/188) showed a significant decrease (Figure 6C and Figure 6−figure supplement 2). Considering that SNr neurons receive inputs from the GPe as well as other brain regions (Foster et al., 2021; Karube et al., 2019; Lee et al., 2020; Pamukcu et al., 2020), we focused subsequent analyses on neurons with significant changes in firing rate following GPe PV inactivation. SNr neurons showing increased firing are likely to receive direct inhibitory input from GPe PV neurons, whereas those with decreased firing may reflect indirect network effects mediated by local or long-range projections. By comparing full and reduced GLM models at the single-neuron level, we found that for each of five task events (Go, No-Go, lick-bout onset, Hit, and CR), the distributions of neurons with significant positive, significant negative, and non-significant encoding differed significantly between firing-increased and firing-decreased SNr neurons (p < 0.01, χ^2^ test, Figure 6−figure supplement 3). These differences suggest that direct versus indirect GPe PV input may be associated with distinct, subpopulation-specific encoding patterns in SNr.

**Figure 6.**
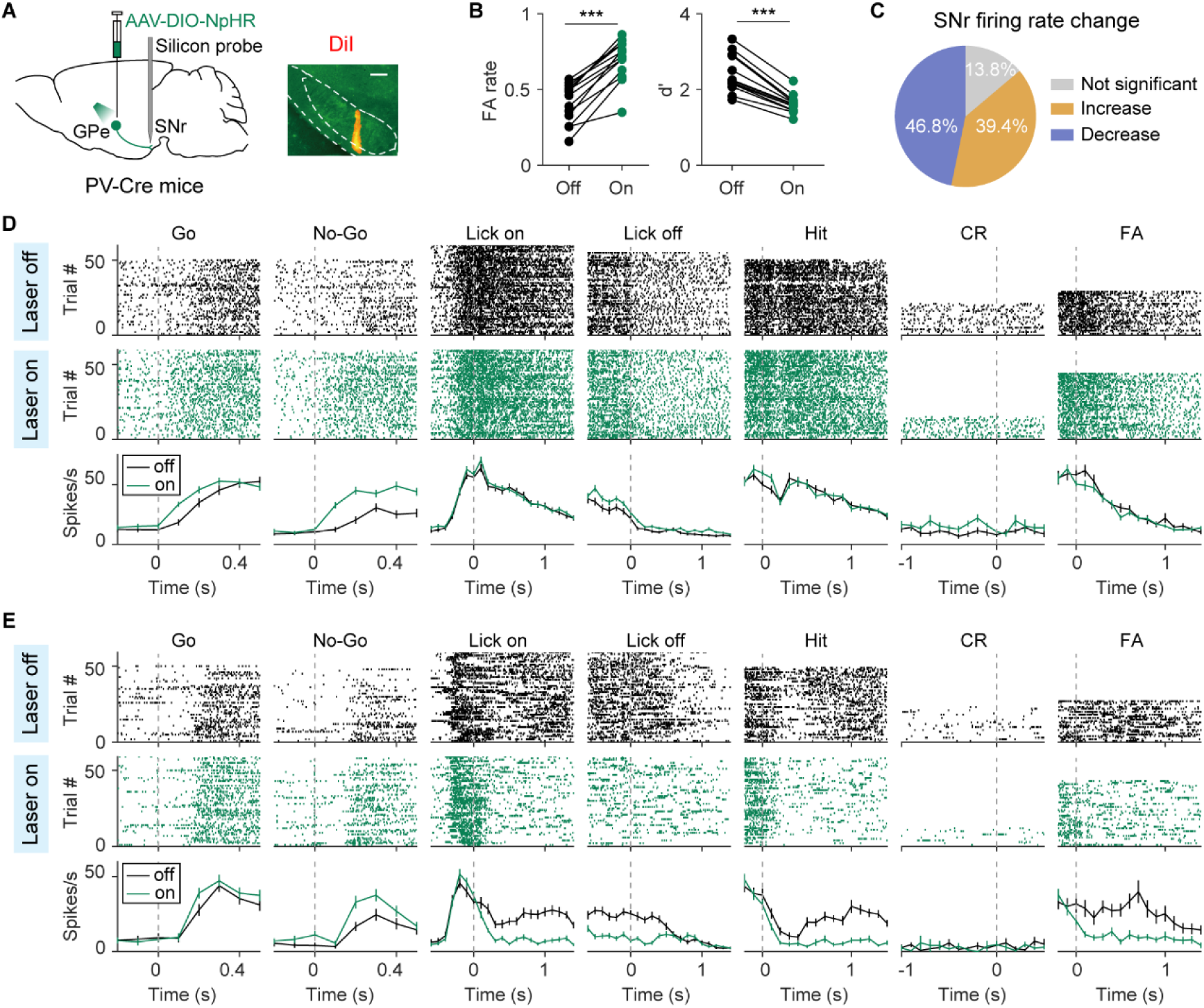
SNr neuronal activity during GPe PV neuron inactivation in task performance. (A) Left, schematic of SNr recording combined with GPe PV neuron inactivation. Right, example fluorescence image showing electrode track labeled with DiI in SNr. Scale bar, 200 μm. (B) In PV-Cre mice used for SNr recordings, inactivation of GPe PV neurons significantly increased FA rate and decreased d’. n = 13 sessions. *** p < 0.001, Wilcoxon signed-rank test. (C) Pie chart showing proportions of SNr neurons (n = 188) with significantly increased, decreased, or unchanged firing rates after inactivation of GPe PV neurons. (D) Spike rasters and PSTHs of an example SNr neuron with a significant increase in mean firing rate after inactivation of GPe PV neurons. For display purposes, 50 − 60 Go trials and 50 − 60 No-Go trials were shown. (E) Same as (D), but for an example SNr neuron with a significant decrease in mean firing rate following GPe PV neuron inactivation. Data represent mean ± SEM.

To further dissect how GPe PV inhibition affects the temporal dynamics of task-related encoding in SNr neurons (two example neurons are shown in Figure 6D and E), we analyzed temporal-kernel GLM coefficients under laser-off and laser-on conditions. This analysis was performed separately for SNr neurons whose firing rates increased or decreased following GPe PV neuron inactivation.

Among putative GABAergic SNr neurons with increased firing rate during GPe PV inactivation, both the temporal profiles of GLM coefficients and the mean coefficients within specific time windows revealed a significant increase for the No-Go stimulus (F_(1, 73)_ = 23.43, p = 7.04×10^-6^, two-way repeated measures ANOVA, Figure 7A; p = 4.93×10^-5^, two-way repeated measures ANOVA followed by Sidak’s multiple comparisons test, Figure 7B) and a significant decrease for lick-bout offset (F_(1, 73)_ = 9.02, p = 0.0036, two-way repeated measures ANOVA, Figure 7A; p = 0.025, two-way repeated measures ANOVA followed by Sidak’s multiple comparisons test, Figure 7B). Similar to the analyses performed for GPe neurons, we also computed the cue SI and outcome SI for these SNr neurons. Cue SI was significantly reduced, both when computed from the average coefficients in the 0 – 0.5 s window after stimulus onset (p = 9.6×10^-7^, n = 74, Wilcoxon signed-rank test, Figure 7C, left) and using sliding-window analysis over time (F_(1, 73)_ = 52.37, p = 3.79×10^-10^, two-way repeated measures ANOVA, Figure 7C, right). This indicates that inactivation of GPe PV neurons reduced the preference of these SNr neurons for the Go stimulus. In contrast, outcome SI, computed from the average GLM coefficients in the 0 – 1 s window, was not significantly affected by laser stimulation (Figure 7D). While the difference in outcome SI between laser-on and laser-off conditions was statistically significant in the sliding-window analysis (F_(1, 73)_ = 4.67, p = 0.034, two-way repeated measures ANOVA, Figure 7D, right), the magnitude of this effect was small.

**Figure 7.**
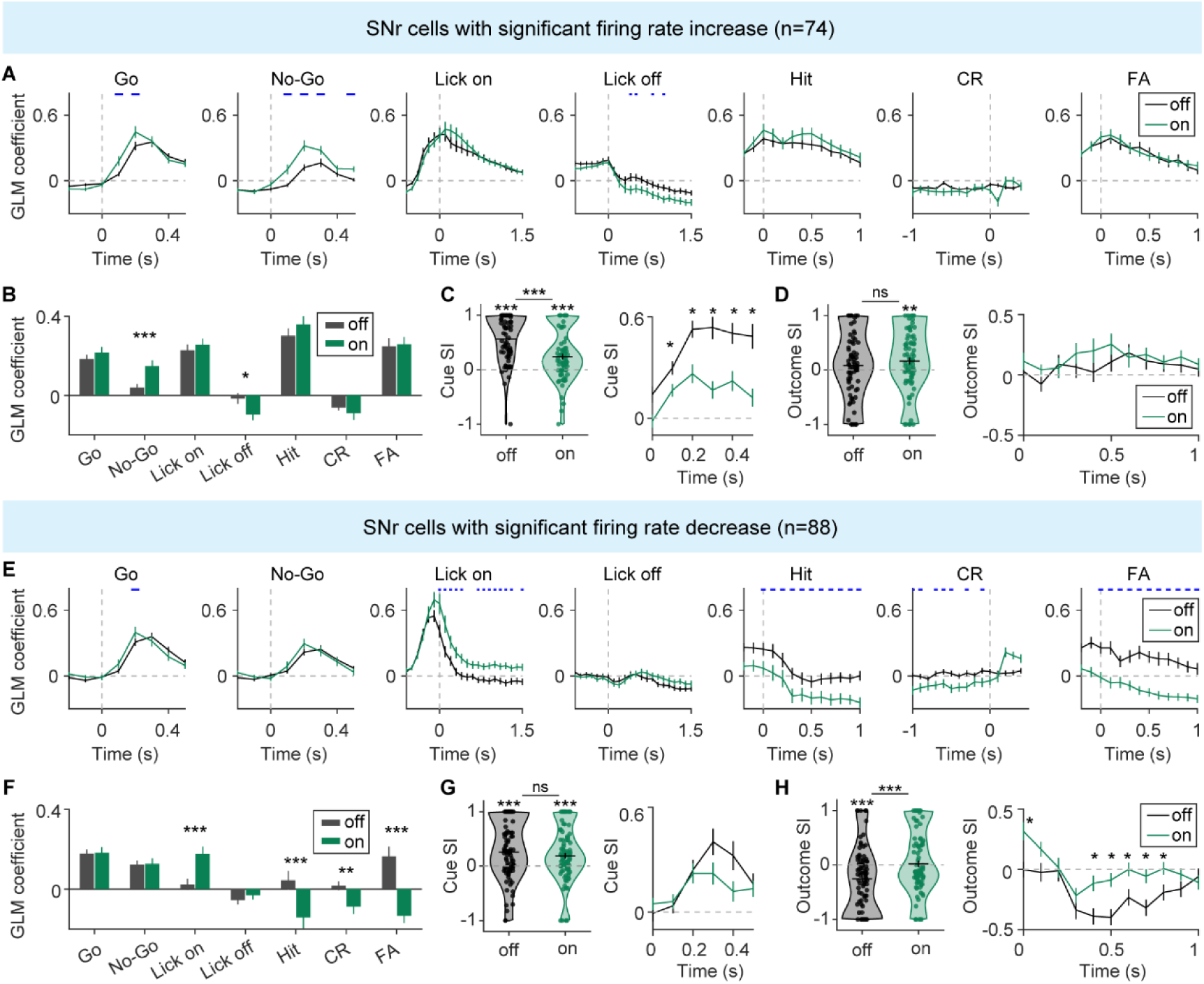
Effect of GPe PV neuron inactivation on task-related encoding of SNr neurons in behaving mice. (A−D) SNr neurons with significantly increased firing rate (n = 74). (A) Temporal profiles of GLM coefficients of each event under laser-off (black) and laser-on (green) conditions. Horizontal ticks indicate time points where GLM coefficients differed significantly between laser-off and laser-on conditions (p < 0.05), determined by a two-way repeated measures ANOVA followed by Sidak’s multiple comparisons test. For each regressor, the two-way ANOVA was applied only to time points within specified time windows: Go or No-Go stimulus, 0 – 0.5 s; lick-bout onset, 0 – 1.5 s; lick-bout offset, 0 – 1.5 s; Hit, 0 – 1 s; CR, −1 – 0 s (with time zero defined as the end of the answer period); and FA, 0 – 1 s. (B) GLM coefficients averaged over the same time windows specified in (A). * p < 0.05, *** p < 0.001, two-way repeated measures ANOVA (F_(3.06, 223.1)_ = 4.94, p_interaction_ = 0.0023) followed by Sidak’s multiple comparisons test. (C) Left, cue SI computed using GLM coefficients (Go and No-Go) averaged over 0 − 0.5 s. *** p < 0.001, Wilcoxon signed-rank test. Right, temporal dynamics of cue SI. * p < 0.05, two-way repeated measures ANOVA (F_(1,73)_ = 52.37, p = 3.79×10^-10^) followed by Sidak’s multiple comparisons test. (D) Left, outcome SI computed using GLM coefficients (Hit and FA) averaged over 0 − 1 s. ** p < 0.01, Wilcoxon signed-rank test. Off vs On: p = 0.051, Wilcoxon signed-rank test. Right, temporal dynamics of outcome SI. F_(1, 73)_ = 4.67, p = 0.034, two-way repeated measures ANOVA. (E−H) SNr neurons with significantly decreased firing rate (n = 88). (E) Temporal profiles of GLM coefficients of each event under laser-off and laser-on conditions, as described in (A). (F) GLM coefficients averaged over the same time windows specified in (A). ** p < 0.01, *** p < 0.001, two-way repeated measures ANOVA (F_(1,87)_ = 15.42, p = 1.72×10^-4^) followed by Sidak’s multiple comparisons test. (G) Left, cue SI computed using GLM coefficients (Go and No-Go) averaged over 0 − 0.5 s. *** p < 0.001, Wilcoxon signed-rank test. Off vs On: p = 0.098, Wilcoxon signed-rank test. Right, temporal dynamics of cue SI. F_(4.32,375.9)_ = 3.28, p_interaction_ = 0.0096, two-way repeated measures ANOVA. (H) Left, outcome SI computed using GLM coefficients (Hit and FA) averaged over 0 − 1 s. *** p < 0.001, Wilcoxon signed-rank test. Right, temporal dynamics of outcome SI. * p < 0.05, two-way repeated measures ANOVA (F_(1,87)_ = 25.79, p = 2.15×10^-6^) followed by Sidak’s multiple comparisons test. Data represent mean ± SEM.

For putative GABAergic SNr neurons with decreased firing rate during GPe PV inactivation, stimulus encoding remained unchanged (Figure 7E and F), and cue SI showed no significant difference between laser-off and laser-on conditions when averaged over the 0 – 0.5 s window, with only modest temporal change (Figure 7G). In contrast, the coefficient for lick-bout onset increased significantly (F_(1, 87)_ = 16.08, p = 1.28×10^-4^, two-way repeated measures ANOVA, Figure 7E; p = 8.95×10^-4^, two-way repeated measures ANOVA followed by Sidak’s multiple comparisons test, Figure 7F), while the lick-bout offset coefficient remained unaffected (Figure 7E and F). Notably, outcome-related coefficients for Hit, CR, and FA trials all decreased significantly under laser-on condition (p < 0.01, two-way repeated measures ANOVA, Figure 7E and F). Outcome SI, computed from the average GLM coefficients in the 0 – 1 s window after outcome onset, was significantly negative during laser-off (p = 9.40×10^-5^, Wilcoxon signed-rank test, Figure 7H, left) and shifted toward zero under laser-on, with a significant difference between conditions (p = 2.90×10^-6^, Wilcoxon signed-rank test, Figure 7H, left). Sliding-window analysis further revealed that laser stimulation significantly shifted outcome SI from negative toward positive values (F_(1,87)_ = 25.79, p = 2.15×10^-6^, two-way repeated measures ANOVA, Figure 7H, right). These results indicate that, under normal conditions, these SNr neurons preferentially encoded the unrewarded (FA) outcome, and that GPe PV inactivation reduced this preference.

Together, these findings indicate that GPe PV neurons exert subpopulation-specific modulation of SNr activity. In putative GABAergic SNr neurons with increased firing during PV inactivation, preference for Go stimulus was significantly reduced, whereas outcome preference remained largely unchanged. By contrast, neurons with decreased firing showed no change in stimulus preference but exhibited a significant reduction in preference for unrewarded outcome.

## Discussion

Using a Go/No-Go visual discrimination task, we found that perturbing GPe PV neuron activity during the stimulus presentation period, either through activation or inactivation, disrupted behavioral performance, whereas manipulations of FoxP2 neurons produced distinct effects, with activation increasing FA rate and inactivation having no measurable impact. Both PV and FoxP2 neurons encoded task-relevant sensory, motor, and outcome signals, yet their activity patterns and encoding properties differed. Notably, inactivation of GPe PV neurons disrupted stimulus and outcome representations in putative GABAergic SNr neurons, revealing a downstream impact on basal ganglia output signaling. Together, these findings indicate that PV and FoxP2 GPe neurons contribute differently to sensorimotor processing.

### Contribution of GPe specific cell types to behavioral performance

Previous studies have shown that activation of GPe PV neurons enhanced movement in healthy mice and restored mobility in PD mouse model (Cui et al., 2021b; Mastro et al., 2017; Pamukcu et al., 2020; Wu and Ding, 2017), while their inactivation did not affect movement under dopamine-depleted conditions (Mastro et al., 2017). Moreover, selective activation and inhibition of GPe PV neurons projecting to the SNr increased and decreased locomotor activity, respectively (Lilascharoen et al., 2021). Activating GPe FoxP2 or Npas1 neurons, on the other hand, suppressed locomotion (Aristieta et al., 2021; Labouesse et al., 2023; Pamukcu et al., 2020).

In the present study, we used both gain- and loss-of-function approaches to dissect the roles of GPe PV and FoxP2 neurons in perceptual decision-making. We found that both activation and inactivation of GPe PV neurons during the stimulus presentation period impaired behavior, which may involve downstream circuits such as SNr neurons. Consistent with this, our recordings during GPe PV inactivation revealed subpopulation-specific changes in SNr neuronal encoding. In contrast, only activation of FoxP2 neurons affected behavior, likely through suppression of striatal activity, which itself is causally involved in perceptual decision-making (Cox and Witten, 2019; Cui et al., 2025; Ding, 2023; Ding and Gold, 2012; Liu et al., 2023; Sippy et al., 2015; Yartsev et al., 2018). Importantly, manipulating either PV or FoxP2 neurons during ITI did not produce significant increases in licking behavior compared to controls, suggesting that the behavioral impairments were not due to changes in motor output per se, but rather a disruption of task-relevant sensory or decision-related processing. These findings support the growing view that GPe neurons contribute to both motor and non-motor functions (Isett et al., 2023; Schechtman et al., 2016; Yuan et al., 2017). Notably, GPe PV neurons consist of distinct subpopulations projecting to the STN/SNr or PF (Courtney et al., 2023; Lilascharoen et al., 2021), though whether each contributes to task performance remains to be determined.

Previous studies showed that inhibiting GPe Npas1 neurons increased movement (Pamukcu et al., 2020), and that GPe arkypallidal neurons were active in canceling imminent action during a stop-signal task (Mallet et al., 2016). However, in our Go/No-Go task, inactivation of GPe FoxP2 neurons—a subpopulation of Npas1 neurons—did not alter behavioral performance. These results suggest that the effect of inhibiting GPe FoxP2 neurons on motor or cognitive control may depend on behavioral context, with minimal impact in tasks that primarily rely on sensory-guided decision-making.

### Task-relevant encoding of GPe PV and FoxP2 neurons

In vivo whole-cell recordings from anesthetized mice revealed that arkypallidal and prototypic GPe neurons displayed distinct multiphasic response patterns to whisker stimulation via air puff (Johansson and Ketzef, 2023; Ketzef and Silberberg, 2021). In rats performing a novel environment exposure task, population activity of GPe neurons encoded environmental identity (Peer et al., 2022), suggesting that specific GPe neurons process sensory features of the surroundings. Building on these studies, our GLM analysis showed that both PV and FoxP2 neurons in the GPe encoded visual stimuli, with PV neurons containing a larger fraction of cells exhibiting significantly positive encoding of the Go stimulus. Moreover, PV neurons showed stronger preference for Go stimulus during the early phase of visual stimulation. Given that the Go stimulus reliably signaled reward while the No-Go stimulus did not, this preference suggests that PV neurons preferentially encode reward-associated sensory cues. These observations are consistent with findings in rats, where faster-firing GPe PV neurons were more strongly modulated by value than arkypallidal neurons (Farries et al., 2023), and with primate studies reporting that high-frequency discharge GPe neurons exhibited selective responses to reward-predicting cues (Joshua et al., 2009; Katabi et al., 2023).

Fiber photometry recordings showed that GPe PV and Npas1 neurons increased activity at locomotion onset (Labouesse et al., 2023; Lilascharoen et al., 2021). In vivo extracellular recordings from moving mice revealed heterogeneous activity patterns among prototypic GPe neurons during movement, whereas FoxP2-expressing arkypallidal GPe neurons exhibited firing rate increases at movement onset, with both cell types reliably encoding movement (Dodson et al., 2015). Extending these observations to a goal-directed behavioral context, we found that both PV and FoxP2 neurons encoded licking behavior in a Go/No-Go task, with a large fraction of neurons in both populations showing positive encoding of lick-bout onset. Notably, the mean GLM coefficients of PV neurons at lick-bout offset were significantly more negative than those of FoxP2 neurons, suggesting a differential representation of action termination between the two cell types.

Beyond sensory and motor processing, GPe neurons are also implicated in encoding reward-related information, including reward outcome, prediction, and value (Arkadir et al., 2004; Farries et al., 2023; Joshua et al., 2009; Katabi et al., 2023; Kim et al., 2017; Nougaret and Ravel, 2018; Yoshida and Hikosaka, 2025). Consistent with findings in monkey demonstrating GPe neuron responses to reward outcomes (Joshua et al., 2009), we observed that both PV and FoxP2 GPe neurons contained subpopulations with positive or negative outcome encoding, with FoxP2 neurons including a larger fraction of cells showing significant encoding of the unrewarded outcome. PV neurons, on the other hand, exhibited a stronger preference for rewarded over unrewarded outcome during early post-outcome period. The mechanisms underlying these cell type-specific differences in reward-related activity and their functional implications remain to be explored.

Retrograde tracing studies have shown that GPe neurons receive inputs from diverse brain regions, including the striatum, STN, thalamus, cortex, amygdala, hypothalamus, and midbrain regions (Abecassis et al., 2020; Chang et al., 2022; Cui et al., 2021a; Ferenczi et al., 2025; Hunt et al., 2018; Lilascharoen et al., 2021; Ma et al., 2025; Shen et al., 2024; Tian et al., 2024). The task-related activity observed in GPe PV and FoxP2 neurons likely reflects integration of these diverse inputs. For instance, stimulus encoding may arise from cortical or thalamic input (e.g., secondary motor cortex and parafascicular nucleus) (Mandelbaum et al., 2019; Zatka-Haas et al., 2021), motor-related activity from the STN, striatum, or motor cortex (Magno et al., 2019), and outcome responses from midbrain signals (Fan et al., 2012). These possibilities highlight the GPe as a key node for integrating sensory, motor, and reward signals to support goal-directed behavior. The different task-related signals observed in PV and FoxP2 neurons could reflect their distinct patterns of cortical and subcortical inputs (Ma et al., 2025). Further studies are required to delineate how each input contributes to the dynamic activity of these GPe subpopulations during behavior.

### GPe modulation of SNr activity

A recent study demonstrated that optogenetic stimulation of GPe PV neurons caused heterogeneous changes in spontaneous firing rates of SNr neurons, with a large fraction exhibiting reduced firing rates and a small subset showing increased activity (Aristieta et al., 2024). In dopamine-depleted mice, optogenetic stimulation of GPe PV neurons attenuated pathological activity in the SNr by reducing the proportion of bursting neurons (Mastro et al., 2017). In monkeys performing a sequential saccade choice task, blocking GABAergic inputs to GPe with bicuculine impaired the suppression of saccades to low-value objects and reduced the excitatory responses of SNr neurons to these objects, without affecting SNr responses to high-value objects (Amita and Hikosaka, 2019). However, it remains unclear how GPe PV and FoxP2 neurons differentially influence SNr activity. Although GPe FoxP2 neurons primarily project to the striatum (Abdi et al., 2015), they may influence SNr activity indirectly via polysynaptic pathways. We found that inactivation of GPe FoxP2 neurons in awake, non-behaving mic had only modest effects on the spontaneous activity of putative GABAergic SNr neurons, suggesting that FoxP2 neurons are not a major determinant of SNr firing under resting or non-task conditions. In contrast, GPe PV neurons inactivation induced significant bidirectional changes in spontaneous firing rates in more than half of recorded SNr neurons. During the Go/No-Go visual task, optogenetic inactivation of GPe PV neurons again induced bidirectional changes in firing rates of two SNr subpopulations, modulating responses related to stimuli, licking, and outcomes. The observed decreases in firing rate may reflect intra-SNr inhibitory interactions via axon collaterals (Brown et al., 2014; Deniau et al., 1982; Deniau et al., 2007). Such heterogeneous changes in SNr firing rates align with previous findings that manipulating striatal D1- or D2-MSNs can produce both increases and decreases in SNr firing (Chen et al., 2021; Freeze et al., 2013). Notably, inactivation of GPe PV neurons reduced the preference for the Go stimulus in the subpopulation of SNr neurons with increased firing and attenuated the preference for unrewarded outcome in those with decreased firing. These results suggest that under normal conditions, GPe PV neurons shape stimulus- and outcome-related encoding in distinct SNr subpopulations, supporting precise modulation of task-relevant signals in the basal ganglia output pathway.

### Limitations of the study

The GPe PV neurons identified in our study likely included heterogeneous subtypes projecting to distinct downstream targets, including the STN, SNr, and PF (Courtney et al., 2023; Lilascharoen et al., 2021). Further work is needed to dissect the specific sensorimotor roles of these PV neuron subtypes during perceptual decision-making. In addition, although we identified two SNr neuron populations with distinct task-related encoding that were differentially modulated by GPe PV neuron inactivation, their respective contributions to behavior remain to be determined.

## Materials and methods

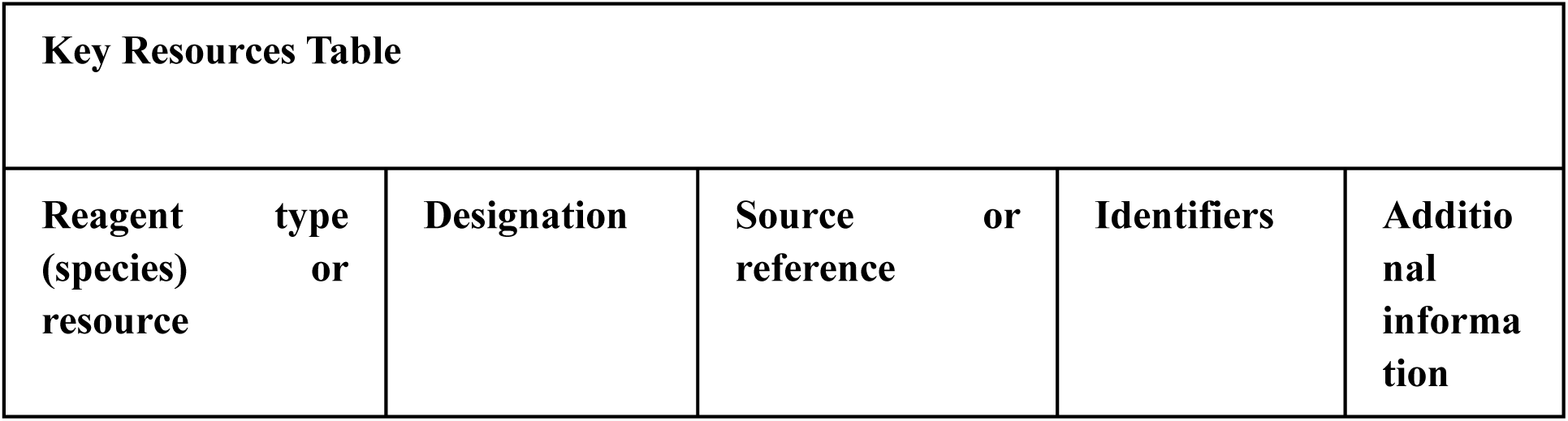
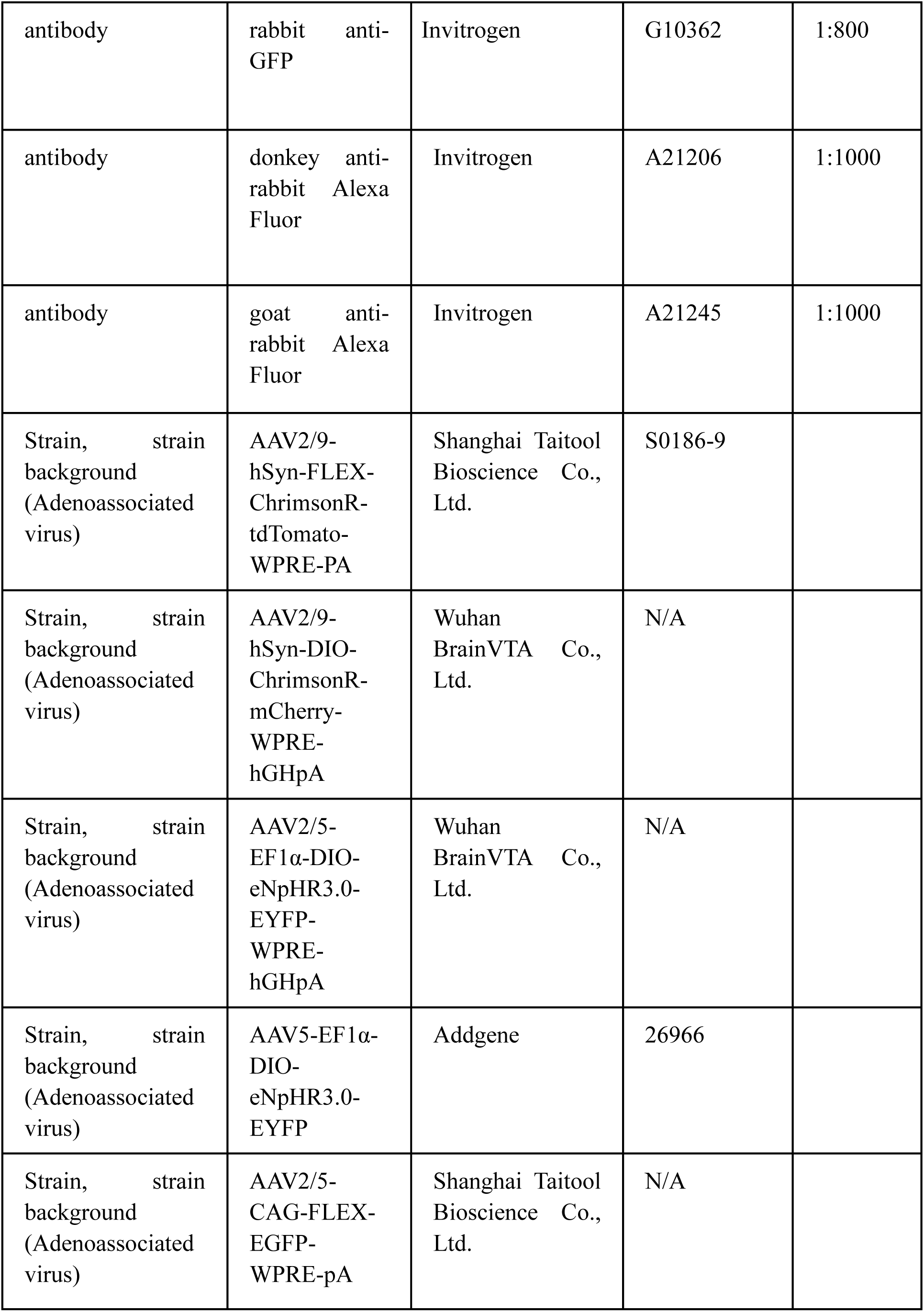
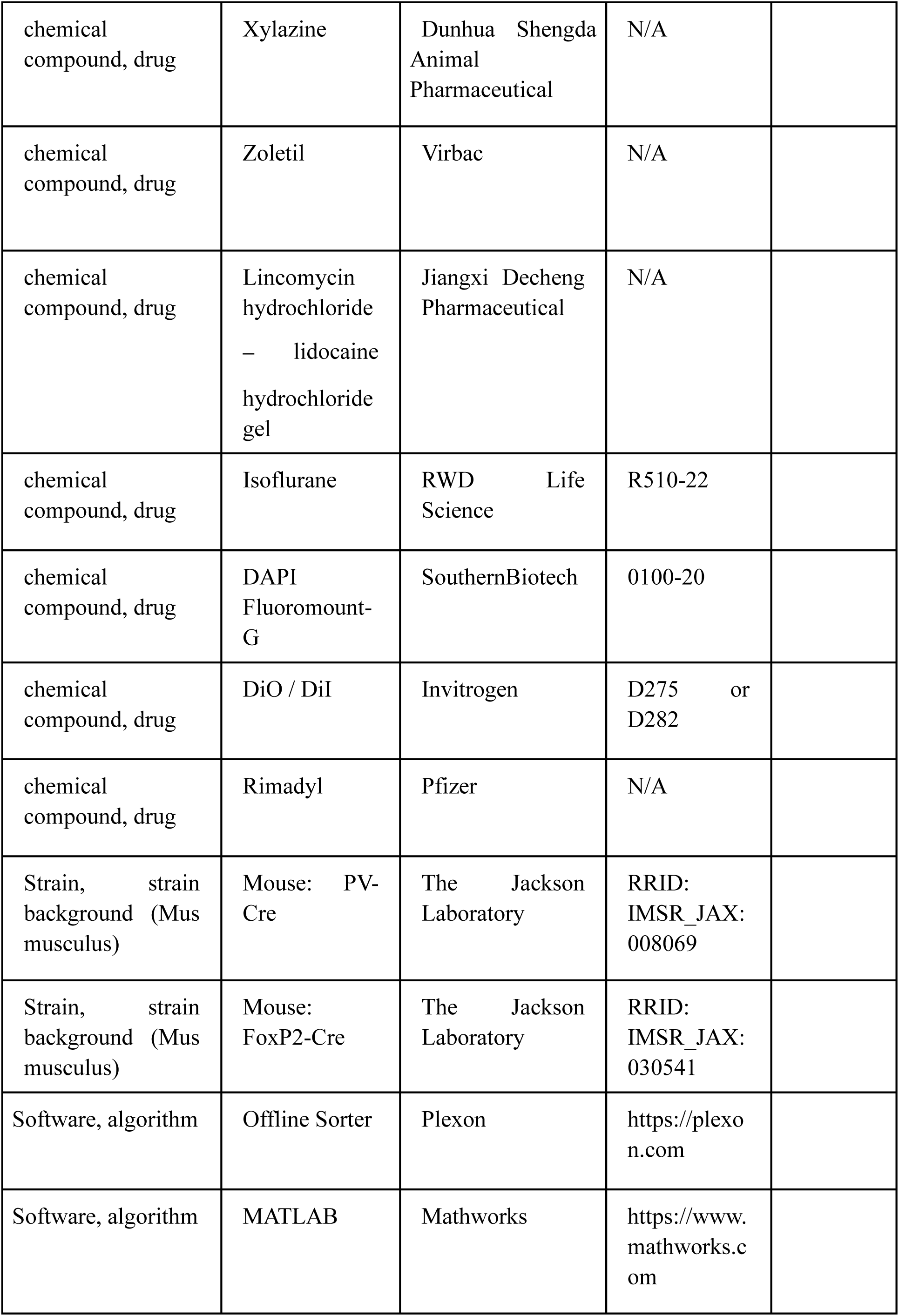
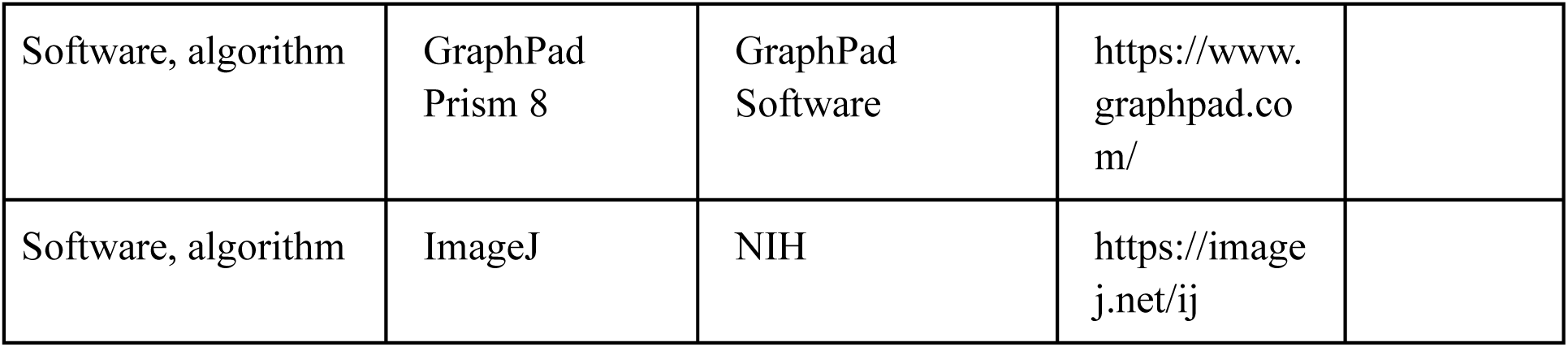

### Animals

All animal procedures (IACUC No. NA-013-2025) were approved by the Animal Care and Use Committee at the Center for Excellence in Brain Science and Intelligence Technology (the Institute of Neuroscience), Chinese Academy of Sciences, and were conducted in accordance with the Center’s guidelines for the care and use of laboratory animals. PV-Cre (The Jackson Laboratory, Bar Harbor, ME, USA, stock No. 008069) and FoxP2-Cre (The Jackson Laboratory, Bar Harbor, ME, USA, stock No. 030541) mice were used in the experiments. Adult (2 – 4 months old) mice of both sexes were used for the experiments. Mice were housed under standardized conditions (temperature: 22 – 23 °C, humidity: 40 – 70 %, 12-hour light/12-hour dark cycle) in the Center’s animal facility.

### Surgery and virus injection

The mice were anesthetized via intraperitoneal injection of a mixture of xylazine (40 mg/kg) and zoletil (30 mg/kg), then placed in a stereotaxic apparatus. The scalp was shaved, and lincomycin hydrochloride–lidocaine hydrochloride gel was applied to the incision site. A unilateral craniotomy (∼ 0.5 mm diameter) was made above the targeted virus injection site. The virus (3 − 5×10^12^ viral particles/mL) was injected using a glass pipette (10 – 20 μm tip diameter) connected to a syringe pump (Harvard Apparatus, Holliston, MA, USA) at a rate of 100 nL/min. After completion of virus injection, the pipette was left in place for at least 10 min before being removed.

To activate GPe PV or FoxP2 neurons, 100 nL of AAV2/9-hSyn-FLEX-ChrimsonR-tdTomato-WPRE-PA was injected at a depth of 3400 μm into the anterior GPe (AP −0.4 mm, ML 2.13 mm) of PV-Cre or FoxP2-Cre mice. An optic fiber (200 μm diameter, NA 0.5) was implanted 450 μm above the virus injection site in GPe. For inactivation of GPe PV (FoxP2) neurons, 200 (120) nL of AAV2/5-EF1α-DIO-eNpHR3.0-EYFP-WPRE-hGHpA or AAV5-EF1α-DIO-eNpHR3.0-EYFP was injected at a depth of 3250 μm into the GPe of PV-Cre (FoxP2-Cre) mice. An optic fiber (400 μm diameter, NA 0.5) was implanted approximately 350 μm above the virus injection site in GPe. In control PV-Cre (FoxP2-Cre) mice, 200 (120) nL of AAV2/5-CAG-FLEX-EGFP-WPRE-pA was injected into the GPe. For optogenetic tagging of PV or FoxP2 neurons, 200 nL of AAV2/9-hSyn-FLEX-ChrimsonR-tdTomato-WPRE-PA or AAV2/9-hSyn-DIO-ChrimsonR-mCherry-WPRE-hGHpA was injected at a depth of 3400 μm into the GPe of PV-Cre or FoxP2-Cre mice. For behavioral experiments, a stainless-steel headplate was implanted and secured to the skull with dental cement. For extracellular recordings, the skull surface above the GPe (AP −0.4 mm, ML 2.13 mm) or SNr (AP −3.16 mm, ML 1.30 mm) was marked with permanent ink. After surgery, lincomycin hydrochloride–lidocaine hydrochloride gel was applied to the incision site to prevent infection and relieve local pain. Mice were then given Rimadyl via their drinking water for 3 days for systemic analgesia, and were allowed to recover for at least 10 days before behavioral training.

### Behavioral task

Visual stimuli used in the Go/No-Go task were oriented gratings (114° × 103°, contrast = 100%, spatial frequency = 0.04 cycles/deg), which were presented on a 19’ LCD monitor (Dell P1917S, mean luminance 35 − 40 cd/m^2^, refresh rate 60 Hz) placed 12 cm away from the eye contralateral to the virus injection site or the recording site. The Go stimulus (vertical grating) and No-Go stimulus (horizontal grating) were randomly interleaved. Each stimulus presentation period consisted of a waiting period, during which the grating was static, followed by an answer period, during which the vertical grating drifted rightward and the horizontal grating drifted upward.

Before training, mice were water-deprived for 2 days. During training, each mouse was head-fixed in a custom apparatus and placed in an acrylic tube inside the behavioral chamber. A lick spout was placed approximately 2 mm posterior to the mouse’s nose tip and 1 mm below the lower lip. Licks were detected either using a custom-made electrical lick sensor or via the interruption of an infrared beam. Water was delivered through a peristaltic pump (Kamoer, Shanghai, China). An Arduino-based microcontroller system was used for stimulus presentation, lick detection, water delivery, laser stimulation, and data acquisition. Task-related signals and lick signals were sampled at 1000 Hz.

Behavioral training consisted of three phases: habituation, conditioning, and visual discrimination. During the habituation phase (2 days), the mouse learned to lick the lickspout to receive a water reward (4 − 5 μL) delivered every 5 s. In the conditioning phase (2 days), a vertical grating was presented on each trial. Each trial consisted of the stimulus presentation period and an inter-trial interval (ITI), defined as the period between the end of the answer period of the previous trial and the onset of the next stimulus. The ITI began with a 4-s blank screen, followed by a variable blank period. During this period, the mouse was required to withhold licking for 3 s to terminate the trial and initiate the next stimulus. If a lick occurred during this period, the 3-s countdown was reset, extending the interval, until either the mouse successfully withheld licking for 3 s or the maximum duration of 13 s was reached. The presentation of the grating stimulus consisted of a 0.5-s waiting period followed by a 2.5-s answer period. A lick detected during the answer period triggered a water reward (4 − 5 μL). During the visual discrimination phase (7 – 21 days), the Go and No-Go trials were randomly interleaved. The timing of the waiting period, answer period, and ITI was the same as in the conditioning phase. In each trial, licking within the answer period were used to assess behavioral performance. In Go trials, licking within the answer period was rewarded with 4 − 5 μL of water (Hit), while no lick was counted as a Miss. In No-Go trials, licking during the answer period was considered a FA, and no lick was a correct rejection (CR). Each mouse performed the task for 1 − 1.5 h per session. For mice undergoing electrophysiological recording, the first lick in the answer period of a No-Go trial triggered a sound cue (100 ms duration), which served as a temporal marker for the onset of FA events in the GLM analysis.

### Optogenetic stimulation

Optical activation of NpHR was induced by green light, and activation of ChrimsonR was induced by red light. A green (532 nm) or a red (635 nm) laser (Shanghai Laser & Optics Century Co., Shanghai, China) was connected to an output optical fiber. Laser stimulation was controlled by an Arduino microcontroller. For in vivo experiments, laser power measured at the fiber tip was 5 – 20 mW for the green laser and 0.5 – 2 mW for the red laser.

For optogenetic activation (or inhibition) of PV or FoxP2 neurons during task performance, laser-off and laser-on blocks (20 trials/block) were interleaved within each session. When laser stimulation was applied during the stimulus presentation period, the laser was delivered from 0 to 3 s after stimulus onset in both Go and No-Go trials. For stimulation during the ITI of the task, laser was applied for 2 s during 6 – 8 s after stimulus onset, which was within the period between the end of the answer period of the previous trial and the stimulus onset of the next trial.

To assess the effect of inactivating GPe PV or FoxP2 neurons on spontaneous SNr activity, optogenetic stimulation was delivered in 50 cycles of 2-s laser-on and 3-s laser-off.

### Optogenetic tagging

For optogenetic tagging using PV-Cre or FoxP2-Cre mice, red laser stimulation (50 or 100 trials, each lasting 10 or 100 ms, with an inter-trial interval of 7−10 s) was delivered in the GPe through the optotrode fiber. Spike counts in the 1-s period preceding laser onset were compared with those in six post-laser windows, defined as the period from laser onset to 1, 2, 3, 4, 5, or 6 ms after onset, respectively. Each window was tested separately using a paired t-test, and a GPe unit was considered significantly activated at a given latency if p < 0.01. Additionally, the Pearson’s correlation coefficient between the spontaneous spike waveform and the laser-evoked spike waveform had to exceed 0.95 to confirm optogenetic tagging (Chen et al., 2021; Lee et al., 2019).

### In vivo extracellular recording

Mice trained in the Go/No-Go visual discrimination task for at least 7 days were used for extracellular recordings. Recordings were conducted in mice that maintained a d′ greater than 1.5 after 1 h of head-fixation. Prior to recording, mice were placed in an acrylic tube, and their headplates were fixed to a holder mounted on a stereotaxic apparatus. Anesthesia was induced with isoflurane (1–2%), and a craniotomy (∼ 1 mm diameter) was made above the anterior GPe (AP −0.4 mm, ML 2.13 mm) or medial/central SNr (AP −3.16 mm, ML 1.30 mm). After dura removal, the craniotomy was covered with silicone elastomer (Kwik-Cast, WPI, Sarasota, FL, USA). Mice were allowed to recover in their home cage for at least 12 hours before recordings commenced. Recordings were made using optotrodes (ASSY-37-32-1-6mm or ASSY-37-32-1-9mm, Diagnostic Biochips, Glen Burnie, MD, USA). For electrophysiological recordings, we included only sessions in which PV or FoxP2 neurons were identified and found to be task-modulated. Among these, the stimulus contrast was 100% in most sessions (n = 50) and 15% in a subset of sessions (n = 24). Following recording, the electrode was retracted, the craniotomy was cleaned with saline, and covered with silicone elastomer. Each mouse underwent 1 – 6 recording sessions. The electrodes were coated with fluorescent dye (DiO or DiI, Invitrogen, Eugene, OR, USA) to visualize the electrode tracks.

We used a Cerebus 64-channel or 32-channel system (Blackrock Microsystems, Salt Lake City, UT, USA) to amplify and filter neural signals. The signals were band-pass filtered between 250 and 7500 Hz, and the waveforms of spikes were detected by thresholding at 4 standard deviations above background noise. Spiking signals were sampled at 30 kHz. Offline spike sorting was performed using Offline Sorter V3 (Plexon Inc., Dallas, TX, USA) based on cluster analysis of principal component. Spike clusters were classified as single units if the interspike interval exceeded 1 ms and the p-value for multivariate analysis of variance tests on the clusters was less than 0.05.

### Slice recording

We performed whole-cell current-clamp recordings in acute brain slices to verify the efficacy of NpHR in GPe FoxP2 neurons. FoxP2-Cre mice injected with AAV2/5-EF1α-DIO-eNpHR3.0-EYFP-WPRE-hGHpA into the GPe were anesthetized with isoflurane and transcardially perfused with ice-cold, oxygenated (95% O_2_/5% CO_2_) NMDG-HEPES solution containing (in mM): 93 NMDG, 2.5 KCl, 1.2 NaH_2_PO_4_, 30 NaHCO_3_, 20 HEPES, 25 glucose, 5 sodium ascorbate, 2 thiourea, 3 sodium pyruvate, 10 MgSO_4_.7H_2_O, 0.5 CaC_l2_.2H_2_O, and 12 NAC (pH 7.4, adjusted with HCl). The brain was removed, and coronal slices containing the GPe (250 μm) were prepared using a vibratome (VT1200S, Leica, Deer Park, IL, USA) in ice-cold oxygenated NMDG-HEPES solution. Slices were recovered in oxygenated NMDG-HEPES solution at 32 °C for 10 min and then transferred to artificial cerebrospinal fluid (ACSF) containing (in mM): 125 NaCl, 3 KCl, 2 CaCl_2,_ 2 MgSO_4_, 1.25 Na_2_HPO_4_, 1.3 sodium ascorbate, 0.6 sodium pyruvate, 26 NaHCO_3_, 10 Glucose, 10 NAC (pH 7.4, adjusted with 10 M NaOH), where they were incubated at room temperature for 1–3 h. Recordings were performed using a Multiclamp 700B amplifier and Digidata 1550 digitizer (Molecular Devices, San Jose, CA, USA). Whole-cell recordings were obtained at 35 °C in oxygenated ACSF solution containing (in mM): 125 NaCl, 4 KCl, 2 CaCl_2_, 1 MgSO_4_, 1.25 NaH_2_PO_4_, 1.3 sodium ascorbate, 0.6 sodium pyruvate, 26 NaHCO_3_, and 10 glucose (pH 7.4). The electrodes (3 − 5 MΩ) were filled with an internal solution containing (in mM): 135 K-gluconate, 4 KCl, 10 HEPES, 10 sodium phosphocreatine, 4 Mg-ATP, 0.3 Na_3_-GTP (pH 7.2, 276 mOsm). Depolarizing current steps were injected to evoke action potential firing in NpHR-expressing FoxP2 neurons. To activate NpHR, green light (1 s duration, 2 − 6 mW; 13 – 30 sweeps) was delivered through a ×40 objective using an X-Cite LED light source (Lumen Dynamics, Mississauga, Canada). Signals were low-pass filtered at 10 kHz and sampled at 10 kHz.

### Histology

The mice were deeply anesthetized with isoflurane and were perfused with 20 mL of saline followed by 20 mL of 4% paraformaldehyde (PFA). The brains were removed and postfixed in 4% PFA overnight at 4℃. After fixation, the brains were transferred to 30% sucrose in phosphate-buffered saline (PBS) for 2 days at 4℃. The brains were sectioned into 60- or 40-μm coronal slices using a cryostat (NX50, Thermo Fisher Scientific, Waltham, MA, USA). To visualize EYFP fluorescence for brain sections from mice in which AAV2/5-EF1α-DIO-eNpHR3.0-EYFP-WPRE-hGH pA or AAV5-EF1α-DIO-eNpHR3.0-EYFP was injected into the GPe, slices were incubated in blocking solution (20% BSA, 0.5% Triton X-100 in PBS) for 2 h at room temperature, followed by incubation with primary antibodies (rabbit anti-GFP, 1:800, Invitrogen, G10362) overnight at 4℃. After rinsing with PBS, sections were incubated with secondary antibodies (donkey anti-rabbit Alexa Fluor or goat anti-rabbit Alexa Fluor, 1:1000, Invitrogen, A21206 or A21245) for 2 h at room temperature. The sections were then washed in PBS and mounted using DAPI Fluoromount-G (SouthernBiotech, 0100-20, Birmingham, AL, USA); DAPI solution (Solarbio, C0065, Beijing, China) was included in a subset of sections. Fluorescence images were acquired using VS120 (Olympus, Tokyo, Japan). Images were analyzed with ImageJ (NIH, Bethesda, MD, USA).

### Data analysis

Analyses were performed using MATLAB (MathWorks Inc., Natick, MA, USA) or GraphPad Prism (GraphPad Software Inc., San Diego, CA, USA).

Behavioral performance was quantified as follows:

Hit rate = number of Hit trials/(number of Hit trials + number of Miss trials),

FA rate = number of FA trials/(number of FA trials + number of CR trials),

Behavioral discriminability (d′) = norminv(Hit rate) - norminv(FA rate), where norminv is the inverse of the cumulative normal function.

For optogenetic experiments, changes in behavioral performance (ΔHit rate, ΔFA rate, or Δd′) were calculated as the difference between the metric in laser-on and laser-off trials.

To determine whether a neuron was task-modulated, we performed a three-way ANOVA with the following factors: stimulus (Go versus No-Go), action (lick versus no lick), and epoch that contained three periods (a 2-s period during inter-trial interval, the 0.5-s of waiting period, and the first 2-s of answer period). A neuron was considered task-modulated if any main effect or interaction term reached significance (p < 0.01) and was included in the analysis. Only task-modulated neurons with mean firing rate > 1 Hz were included in subsequent analyses.

We identified putative GABAergic SNr neurons based on their spike waveforms (Barter et al., 2015). We computed the peak width, defined as the width at half-maximum of the waveform peak amplitude. Neurons with a peak width less than 0.36 ms were classified as GABAergic SNr neurons (Chen et al., 2021). Only units with baseline firing rates exceeding 5 Hz during the first 10 s of the session were used in subsequent analyses.

To quantify task-related activity of GPe or SNr neurons, we analyzed neuronal responses using a temporal-kernel generalized linear model (GLM) (Pinto and Dan, 2015). Spike counts from each session were binned into 100-ms intervals and Z-scored prior to model fitting. The neuronal response at time *t* was modeled as a linear combination of multiple task events at various temporal delays relative to event onset:

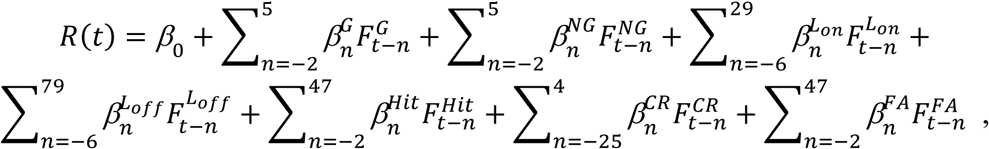

where β_0_ is the offset term, 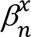 denotes the coefficient for task event *x* at temporal delay *n*, and 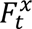 indicates the occurrence of event *x* at time bin *t* (1 if occurred, 0 otherwise). Here, *G* and *NG* represent the Go and No-Go stimuli, respectively; L*_on_* and L*_off_* denote lick-bout onset and offset; and Hit, CR, and FA correspond to reward, correct rejection, and false alarm events, respectively. Lick-bout onset was defined as a lick preceded by at least 2 s without licks; lick-bout offset as the last lick in a bout not followed by another lick within 2 s. The time zero of Hit events was defined as the time of first lick following reward delivery in Go trial, whereas for FA events, it was defined as the time of sound cue triggered by the first lick in the answer period of No-Go trials. The time zero of CR events was defined as the end of answer period, with the analysis focused on neural dynamics related to the decision to withhold licking. Miss trials were not modeled as a separate regressor due to their low occurrence. The model had a total of 268 coefficients plus the offset term. For GPe neurons, only those with more than 10 trials for each event type were included in the GLM analysis. For SNr neurons, only those with more than 5 trials per event type in both laser-off and laser-on conditions were included.

The temporal-kernel GLM models were fit using ridge regression (Huth et al., 2012; Pho et al., 2018; Pinto and Dan, 2015). We set aside 20% of trials as the test dataset. The remaining 80% of trials, used for model fitting, were further split into a training dataset (80%) and a validation dataset (20%). For each neuron, a regularization parameter λ was selected from a range of candidate values by maximizing model performance on the validation dataset. Model performance was quantified by the correlation coefficient (CC) between predicted and measured neural activity in the validation dataset. This cross-validation procedure was repeated 10 times, and the λ value that yielded the highest average CC on the validation sets was chosen. Using this optimal λ, the model was then fit on the combined training and validation data. Final model performance was assessed by computing the CC between the predicted and observed responses in the held-out test dataset. Because spike counts were Z-scored, CC provides a scale-invariant measure of how well the model captures variance in neural activity. Statistical significance was assessed by bootstrapping the test dataset 1000 times to generate a null distribution of CC values. Neurons were considered to have significant GLM fits if the observed CC was positive and exceeded chance level (p < 0.05, bootstrap test). Only neurons with significant GLM fits were included in subsequent analysis.

Temporal profiles of the GLM coefficients captured the time-resolved modulation of neuronal activity for each task event. To quantify event-specific encoding, the mean GLM coefficient for each regressor was computed over specified time windows (Go or No-Go stimulus: 0 – 0.5 s; lick-bout onset: 0 – 1.5 s; lick-bout offset: 0 – 1.5 s; Hit: 0 – 1 s; CR: −1 – 0 s, with time zero defined as the end of answer period; FA: 0 – 1 s). Cue SI was computed as (β_Go_ - β_No-Go_)/(|β_Go_| + |β_No-Go_|), in which β_Go_ and β_No-Go_ represent the average GLM coefficients in the 0 – 0.5 s window after stimulus onset in Go and No-Go trials, respectively. For the sliding-window analysis of the cue SI at each time point, the denominator was fixed as the sum of the mean absolute β_Go_ and the mean absolute β_No-Go_ values computed across all time points. The cue SI ranged from −1 to 1, with negative and positive values indicating preference for No-Go and Go stimuli, respectively. Outcome SI was calculated as (β_Hit_ - β_FA_)/(|β_Hit_| + |β_FA_|), where β_Hit_ and β_FA_ represent the average coefficients in the 0 – 1 s window in Hit and FA trials, respectively. For the sliding-window analysis of the outcome SI at each time point, the denominator was fixed as the sum of the mean absolute β_Hit_ and the mean absolute β_FA_ values computed across all time points. The outcome SI ranged from −1 to 1, with negative values indicating a preference for FA outcome and positive values indicating a preference for Hit outcome.

To efficiently quantify the proportion of neurons that showed significant positive or negative encoding of each task event, we compared full and reduced GLM models in which each task event was represented by a single coefficient, omitting the temporal kernels. The structure of the full model was otherwise identical to the temporal-kernel GLM. For each neuron, the full GLM was fit using an 80:20 training-validation split of the data, and this procedure was repeated 100 times with different random partitions. Elastic-net regularization was applied to prevent overfitting, and the regularization parameter was selected to maximize prediction performance on the validation set. To determine whether a neuron significantly encoded a specific task event, the full model including all events was compared to reduced models in which each event was removed in turn. For each neuron and each event, R^2^ values of the full and reduced models were computed across repetitions, and a one-tailed t-test was used to determine whether removal of the event significantly decreased model performance (p < 0.05). Only neurons with significant task-related encoding in the full model (average R^2^ > 0.01 across 100 repetitions, one-tailed t-test, p < 0.05) were included for classification of their event-specific coefficients as significantly positive or negative, based on coefficient sign and statistical significance.

To determine whether inactivation of GPe PV or FoxP2 neurons significantly affects the spontaneous activity of SNr neurons, firing rates within a 2-s period during and before laser stimulation were compared using the Wilcoxon signed-rank test.

To determine whether inactivation of GPe PV neurons significantly affects the firing rates of SNr neurons during task engagement, we performed a two-way ANOVA on PSTHs (0 ‒ 3 s relative to stimulus onset, 100 ms/bin). Comparisons were made between laser-off and laser-on conditions for both Go and No-Go trials. An SNr neuron was considered significantly affected if the p-value in either Go or No-Go condition was < 0.01. The mean firing rate for each SNr neuron was computed based on spikes within 0 ‒ 3 s relative to stimulus onset.

### Statistics

The statistical analysis was performed in MATLAB (MathWorks Inc., Natick, MA, USA) or GraphPad Prism (GraphPad Software Inc., San Diego, CA, USA). Significance was assessed with Wilcoxon signed-rank test, Wilcoxon rank-sum test, paired t-test, two-sample Kolmogorov-Smirnov test, three-way ANOVA, two-way ANOVA, χ^2^ test, and Fisher’s exact test, as appropriate. Pearson’s correlation was used to compute correlation coefficients. Data were reported as mean ± SEM and p values < 0.05 were considered statistically significant.

## Data availability

The processed data and code used to reproduce the figures are available at Mendeley Data (https://doi.org/10.17632/z6sg8rzhxb.2).

## Acknowledgements

This work was supported by the Brain Science and Brain-like Intelligence Technology - National Science and Technology Major Project (2021ZD0203700/2021ZD0203703), the Strategic Priority Research Program of the Chinese Academy of Sciences (Grant No. XDB1010201), and the National Natural Science Foundation of China (31771151, 32171030, and 32571223 to H.Y., 32100829 to D.L.).

## Author contributions

Xiaotian Pu, Conceptualization, Investigation, Methodology, Data curation, Formal analysis, Visualisation, Writing – review & editing; Jingyu Ren, Methodology, Formal analysis; Taorong Xie, Methodology, Formal analysis; Kexin Zou, Investigation; Jingze Xu, Investigation; Yaping Li, Investigation; Dechen Liu, Conceptualization, Investigation, Methodology, Data curation, Funding acquisition, Writing – review & editing; Haishan Yao, Conceptualization, Resources, Formal analysis, Validation, Supervision, Funding acquisition, Writing – original draft preparation; Writing – review & editing.

## Competing interests

The authors declare no competing interests.

## Figure supplements

**Figure 1 - figure supplement 1.**
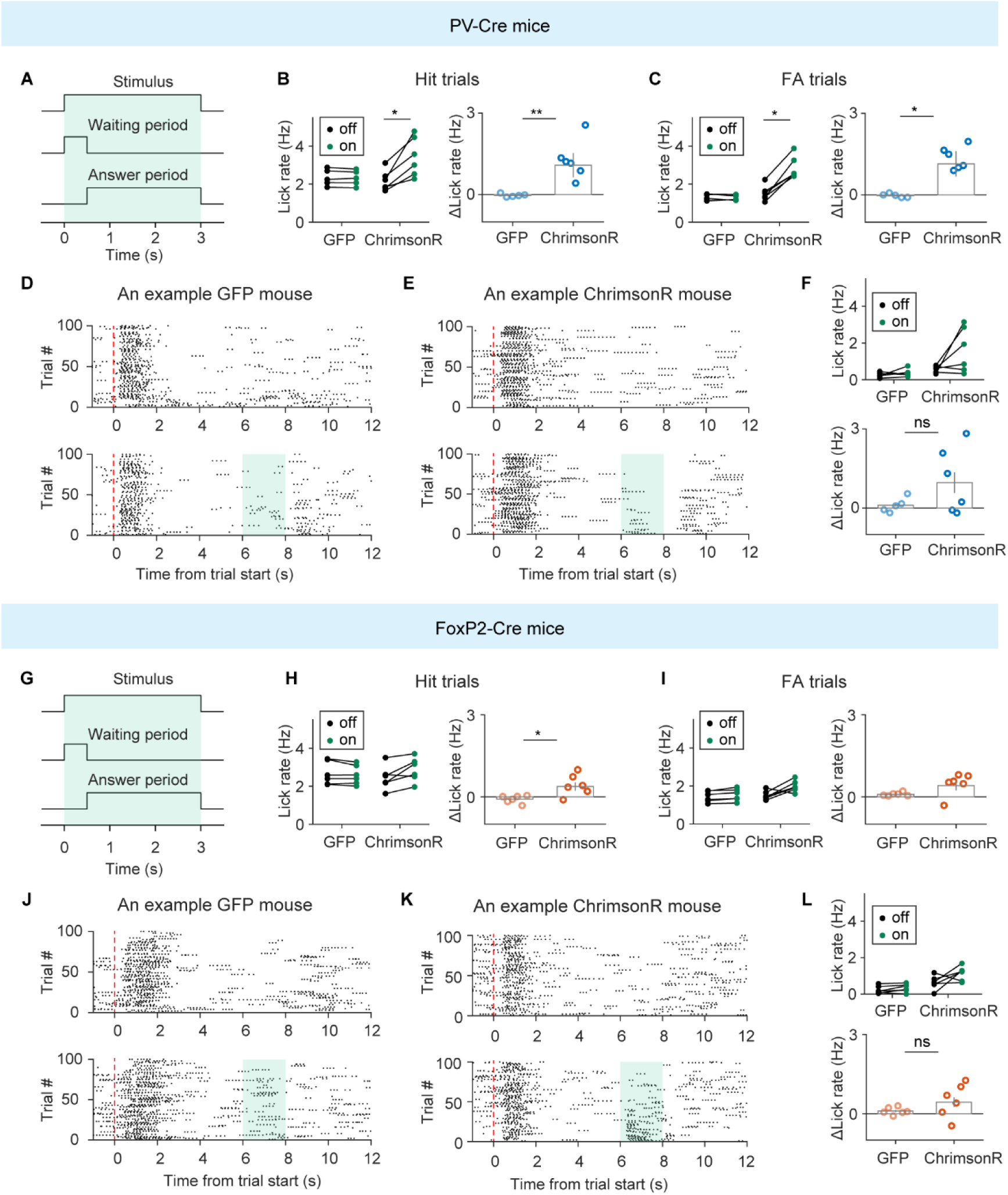
Effect of activating PV or FoxP2 GPe neurons on lick rate during stimulus presentation period or inter-trial interval (ITI). (A−C) Laser stimulation was applied during stimulus presentation period. (A) Schematic of the duration of laser stimulation (green shading) during stimulus presentation period. (B) Left, lick rates during stimulus presentation period in Hit trials under laser-off and laser-on conditions for PV-Cre mice expressing AAV-FLEX-EGFP (n = 5) or AAV-FLEX-ChrimsonR (n = 6) in the GPe. * p < 0.05, Wilcoxon signed-rank test. Right, comparison of lick rate difference between control and ChrimsonR mice. ** p < 0.01, Wilcoxon rank-sum test. (C) Same as (A), but for FA trials. Left, * p < 0.05, Wilcoxon signed-rank test. Right, * p < 0.05, Wilcoxon rank-sum test. (D−F) Laser stimulation was applied during ITI. (D) Lick rasters from an example PV-Cre mouse expressing AAV-FLEX-EGFP in the GPe, shown for ITI laser-off and ITI laser-on trials. Top, laser-off trials. Bottom, laser stimulation was applied during the ITI. Green shading, laser stimulation period. (E) Same as (D), but for a PV-Cre mouse expressing AAV-FLEX-ChrimsonR in the GPe. (F) Top, ITI lick rates under laser-off and laser-on conditions for GFP mice (n = 5, p = 0.63, Wilcoxon signed-rank test) and ChrimsonR PV-Cre mice (n = 6, p = 0.16, Wilcoxon signed-rank test). Bottom, comparison of ITI laser-induced lick rate changes between GFP and ChrimsonR PV-Cre mice (p = 0.28, Wilcoxon rank-sum test). (G−I) Laser stimulation was applied during stimulus presentation period. (G) Schematic of the duration of laser stimulation (green shading) during stimulus presentation period. (H) Left, lick rates during stimulus presentation period in Hit trials under laser-off and laser-on conditions for FoxP2-Cre mice expressing AAV-FLEX-EGFP (n = 6) or AAV-FLEX-ChrimsonR (n = 6) in the GPe. Right, comparison of lick rate difference between control and ChrimsonR mice. * p < 0.05, Wilcoxon rank-sum test. (I) Same as (H), but for FA trials. (J−L) Laser stimulation was applied during ITI. (J) Lick rasters from an example FoxP2-Cre mouse expressing AAV-FLEX-EGFP in the GPe, shown for ITI laser-off and ITI laser-on trials. Top, laser-off trials. Bottom, laser stimulation was applied during the ITI. Green shading, laser stimulation period. (K) Same as (J), but for a FoxP2-Cre mouse expressing AAV-FLEX-ChrimsonR in the GPe. (L) Top, ITI lick rates under laser-off and laser-on conditions for GFP mice (n = 6, p = 0.22, Wilcoxon signed-rank test) and ChrimsonR FoxP2-Cre mice (n = 6, p = 0.16, Wilcoxon signed-rank test). Bottom, comparison of ITI laser-induced lick rate changes between GFP and ChrimsonR FoxP2-Cre mice (p = 0.19, Wilcoxon rank-sum test). For display purposes, 100 laser-off trials and 100 laser-on trials were shown in the raster plot. Data represent mean ± SEM.

**Figure 2 - figure supplement 1.**
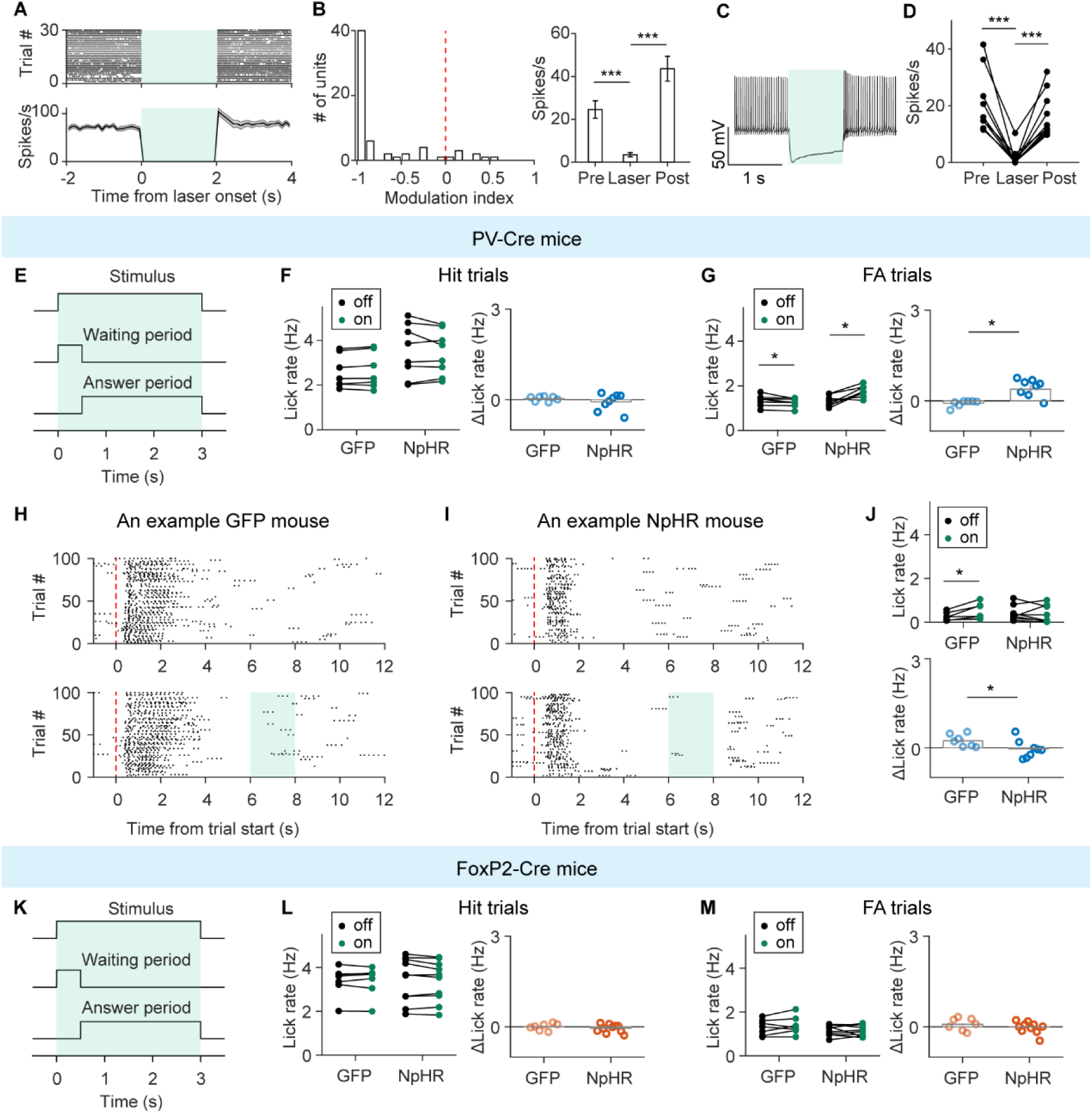
Effect of inactivating PV or FoxP2 GPe neurons on lick rate during stimulus presentation period or inter-trial interval (ITI). (A) Spike raster and PSTH of spontaneous firing from an example GPe neuron recorded in an awake PV-Cre mouse expressing AAV-DIO-NpHR in the GPe. Green shading indicates the laser stimulation period. (B) Optogenetic activation of NpHR significantly suppressed spontaneous firing of GPe neurons in PV-Cre mice expressing AAV-DIO-NpHR in the GPe. Left, distribution of modulation index, defined as (R_laser_ - R_pre_)/(R_laser_ + R_pre_), in which R_laser_ and R_pre_ are the firing rates within a 2-s period during and before laser stimulation, respectively. p = 6.0×10^-11^, n = 64, Wilcoxon signed-rank test. Right, average firing rates before, during, and after laser stimulation. *** p < 0.001, one-way repeated measures ANOVA followed by Sidak’s multiple comparisons test. (C) Responses of an example GPe FoxP2 neuron expressing NpHR, recorded from a brain slice of a FoxP2-Cre mouse, in which AAV-DIO-NpHR was injected into the GPe. Green shading indicates the duration of laser stimulation. (D) Action potentials of NpHR-expressing GPe FoxP2 neurons within a 1-s period before, during and after laser stimulation. *** p < 0.001, one-way repeated measures ANOVA followed by Sidak’s multiple comparisons test. n = 10 cells from 3 FoxP2-Cre mice. (E−G) Laser stimulation was applied during stimulus presentation period. (E) Schematic of the duration of laser stimulation (green shading) during stimulus presentation period. (F) Left, lick rates during stimulus presentation period in Hit trials under laser-off and laser-on conditions for PV-Cre mice expressing AAV-FLEX-EGFP (n = 7) or AAV-DIO-NpHR (n = 8) in the GPe. Right, comparison of lick rate difference between GFP and NpHR mice. p = 0.78, Wilcoxon rank-sum test. (G) Same as (F), but for FA trials. * p < 0.05. Left, Wilcoxon signed-rank test. Right, Wilcoxon rank-sum test. (H−J) Laser stimulation was applied during ITI. (H) Lick rasters from an example PV-Cre mouse expressing AAV-FLEX-EGFP in the GPe, shown for ITI laser-off and ITI laser-on trials. Top, laser-off trials. Bottom, trials with laser stimulation during the ITI. Green shading, laser stimulation period. (I) Same as (H), but for a PV-Cre mouse expressing AAV-DIO-NpHR in the GPe. (J) Top, ITI lick rates under laser-off and laser-on conditions for GFP mice (n = 7, p = 0.016, Wilcoxon signed-rank test) and NpHR PV-Cre mice (n = 8, p = 0.46, Wilcoxon signed-rank test). Bottom, comparison of ITI laser-induced lick rate changes between GFP and NpHR PV-Cre mice (p = 0.025, Wilcoxon rank-sum test). (K−M) Laser stimulation was applied during stimulus presentation period. (K) Schematic of the duration of laser stimulation during stimulus presentation period. (L) Left, lick rates during stimulus presentation period in Hit trials under laser-off and laser-on conditions for FoxP2-Cre mice expressing AAV-FLEX-EGFP (n = 7) or AAV-DIO-NpHR (n = 10) in the GPe. Right, comparison of lick rate difference between GFP and NpHR mice. p = 0.61, Wilcoxon rank-sum test. (M) Same as (L), but for FA trials. Right, comparison of lick rate difference between GFP and NpHR mice. p = 0.34, Wilcoxon rank-sum test. For display purposes, 100 laser-off trials and 100 laser-on trials were shown in the raster plot. Data represent mean ± SEM.

**Figure 3 - figure supplement 1.**
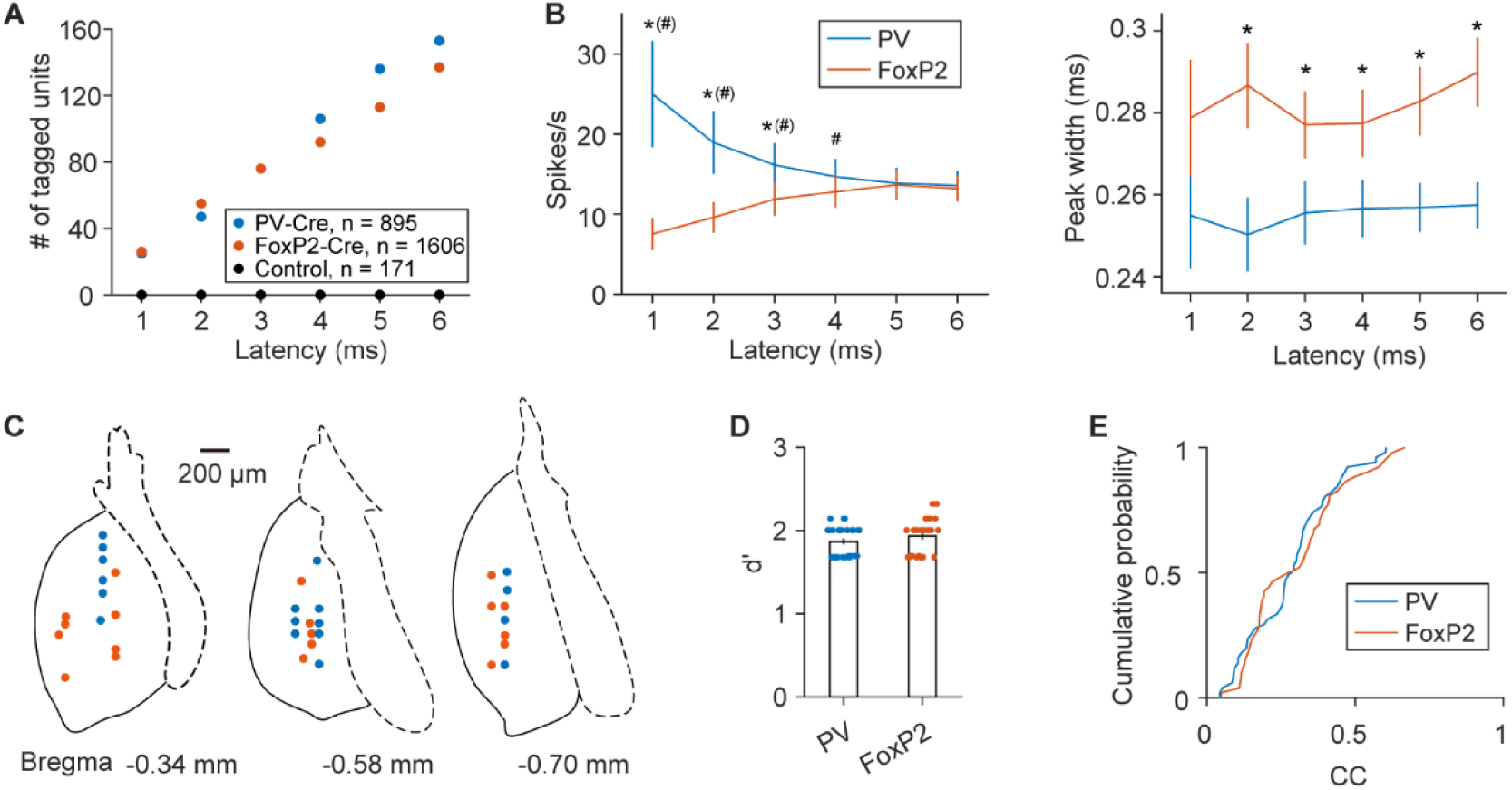
Optogenetic identification, electrophysiological properties, and recording locations of GPe PV and FoxP2 neurons. (A) Number of PV neurons (blue) identified from a total of 895 neurons recorded in PV-Cre mice (50 sessions, 8 mice), and number of FoxP2 GPe neurons (brown) identified from a total of 1606 neurons from FoxP2-Cre mice (85 sessions, 14 mice), across latency thresholds ranging from 1 to 6 ms. In these mice, AAV-FLEX-ChrimsonR was injected into the GPe. In control PV-Cre (4 sessions, 2 mice) and FoxP2-Cre mice (4 sessions, 3 mice) that did not receive virus injection, 72 and 99 neurons were recorded, respectively; none of the 171 neurons met the criteria for optogenetic identification. (B) Left, spontaneous firing rates of PV and FoxP2 neurons identified under different latency thresholds. * p < 0.05, Wilcoxon rank-sum test. # p < 0.05, two-sample Kolmogorov-Smirnov test. Right, spike waveform peak widths of the same populations. * p < 0.05, Wilcoxon rank-sum test. (C) Recording locations of optogenetically identified PV (blue) and FoxP2 (brown) neurons in the anterior GPe. (D) Behavioral performance of PV-Cre mice and FoxP2-Cre mice in which PV and FoxP2 neurons were identified using a 3-ms latency threshold. p = 0.29, Wilcoxon rank-sum test. (E) The correlation coefficients (CCs) between GLM-predicted and actual activity were calculated separately for PV and FoxP2 neurons identified using a 3-ms latency threshold. The distributions of CCs did not differ significantly (p = 0.59, n = 51 and 52 for PV and FoxP2 neurons, respectively, two-sample Kolmogorov-Smirnov test). Data represent mean ± SEM.

**Figure 5 - figure supplement 1.**
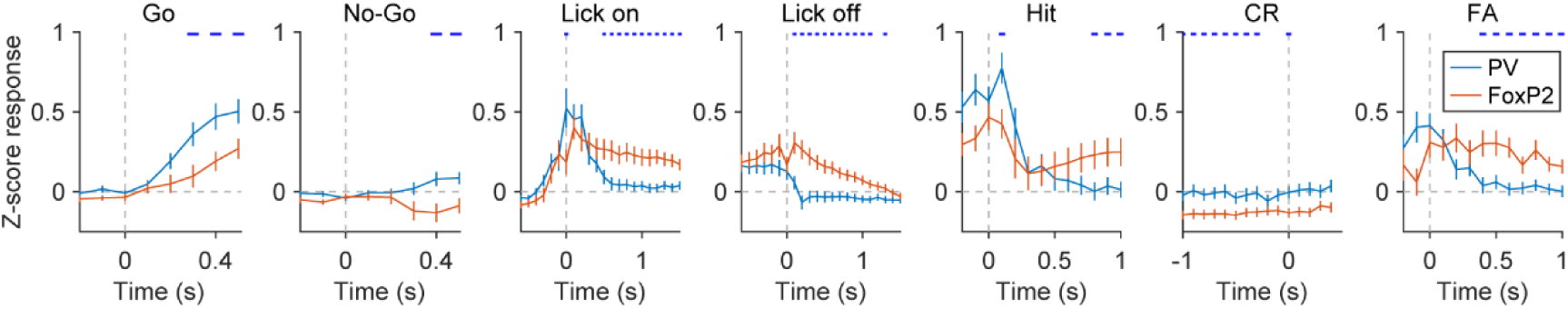
Event-aligned Z-scored responses of task-modulated PV and FoxP2 GPe neurons identified using a 3-ms latency threshold. Average Z-scored neural activity aligned to stimulus onset (in Go and No-Go trials), lick-bout onset, lick-bout offset, and outcome onset (Hit, CR, and FA). Blue, PV neurons (n = 51). Brown, FoxP2 neurons (n = 52). Horizontal ticks mark time points with significant differences between the two cell types (p < 0.05), determined by a two-way ANOVA with mixed design followed by Tukey’s multiple comparisons test. For each event type, the two-way ANOVA was applied only to time points within specified time windows: Go or No-Go stimulus, 0 – 0.5 s; lick-bout onset, 0 – 1.5 s; lick-bout offset, 0 – 1.5 s; Hit, 0 – 1 s; CR, −1 – 0 s (with time zero defined as the end of the answer period); and FA, 0 – 1 s. Data represent mean ± SEM.

**Figure 5 - figure supplement 2.**
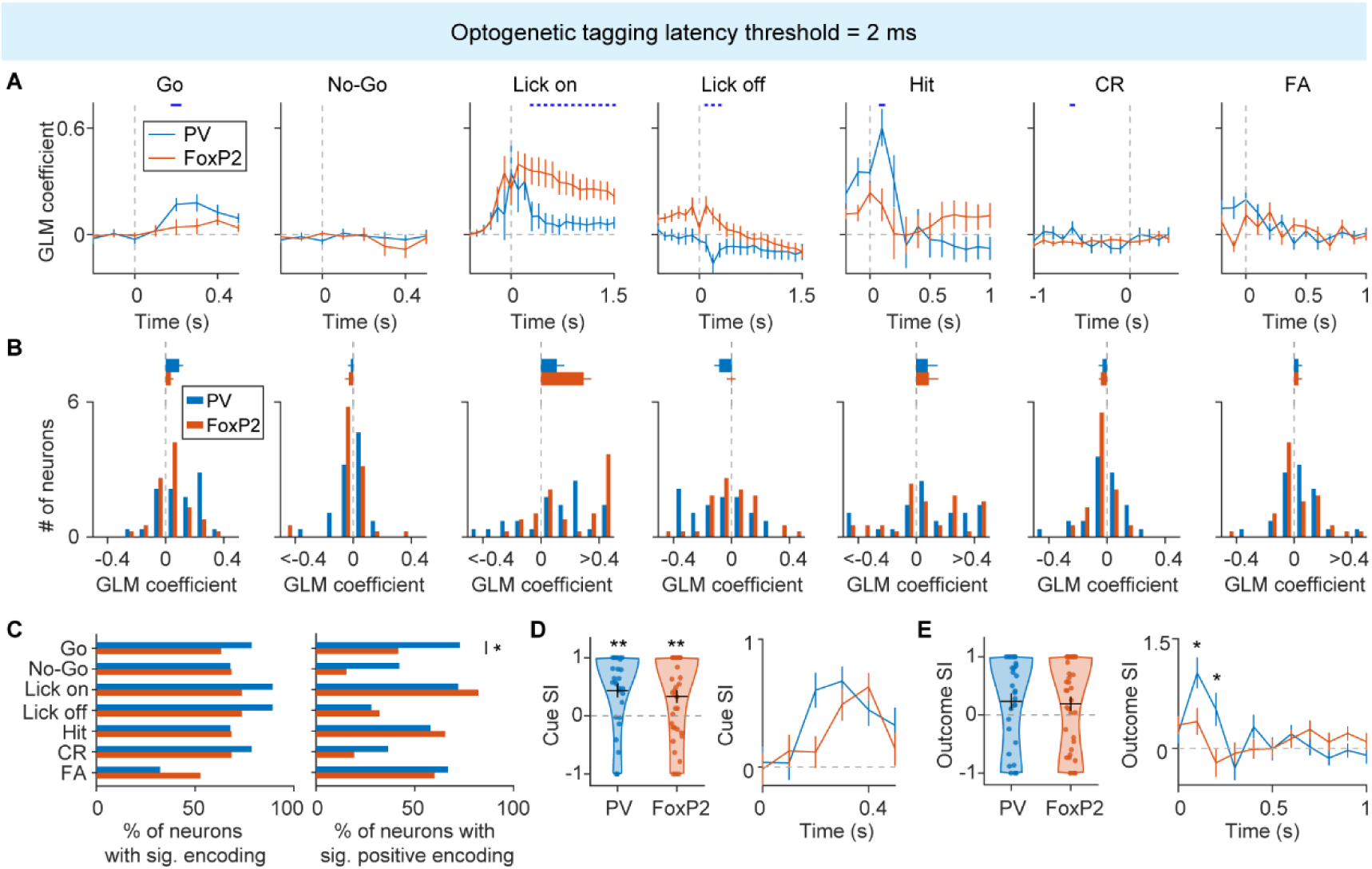
Comparison of GLM coefficients, cue SI, and outcome SI between task-modulated PV and FoxP2 GPe neurons identified using a 2-ms latency threshold. (A) Temporal profiles of GLM coefficient averaged across PV neurons (n = 28) or FoxP2 neurons (n = 38) for each regressor. Horizontal ticks indicate time points where GLM coefficients differed significantly between PV and FoxP2 neurons (p < 0.05), determined by a two-way ANOVA with mixed design followed by Tukey’s multiple comparisons test. For each regressor, the two-way ANOVA was applied only to time points within specified time windows: Go or No-Go stimulus, 0 – 0.5 s; lick-bout onset, 0 – 1.5 s; lick-bout offset, 0 – 1.5 s; Hit, 0 – 1 s; CR, −1 – 0 s (with time zero defined as the end of the answer period); and FA, 0 – 1 s. (B) Distribution of the mean GLM coefficients averaged over the same time windows specified in (A). For Go stimulus, p = 0.035, two-sample Kolmogorov-Smirnov test. Top, average of GLM coefficients. (C) Left, proportion of PV and FoxP2 neurons showing significant encoding for each task event. Right, among neurons with significant encoding, proportion of PV and FoxP2 neurons exhibiting significantly positive (or negative) encoding for each event. * p < 0.05, Fisher’s exact test. (D) Left, cue SI was significantly larger than 0 for both PV and FoxP2 GPe neurons (p < 0.01, Wilcoxon signed-rank test), with no significant difference between the two cell types (p = 0.77, Wilcoxon rank-sum test). Right, temporal dynamics of cue SI in PV versus FoxP2 neurons. (E) Left, outcome SI of PV and FoxP2 neurons. Right, temporal dynamics of outcome SI in PV versus FoxP2 neurons. * p < 0.05, two-way ANOVA with mixed design (F_(10, 640)_ = 4.10, p_interaction_ = 0.0034) followed by Tukey’s multiple comparisons test. Data represent mean ± SEM.

**Figure 5 - figure supplement 3.**
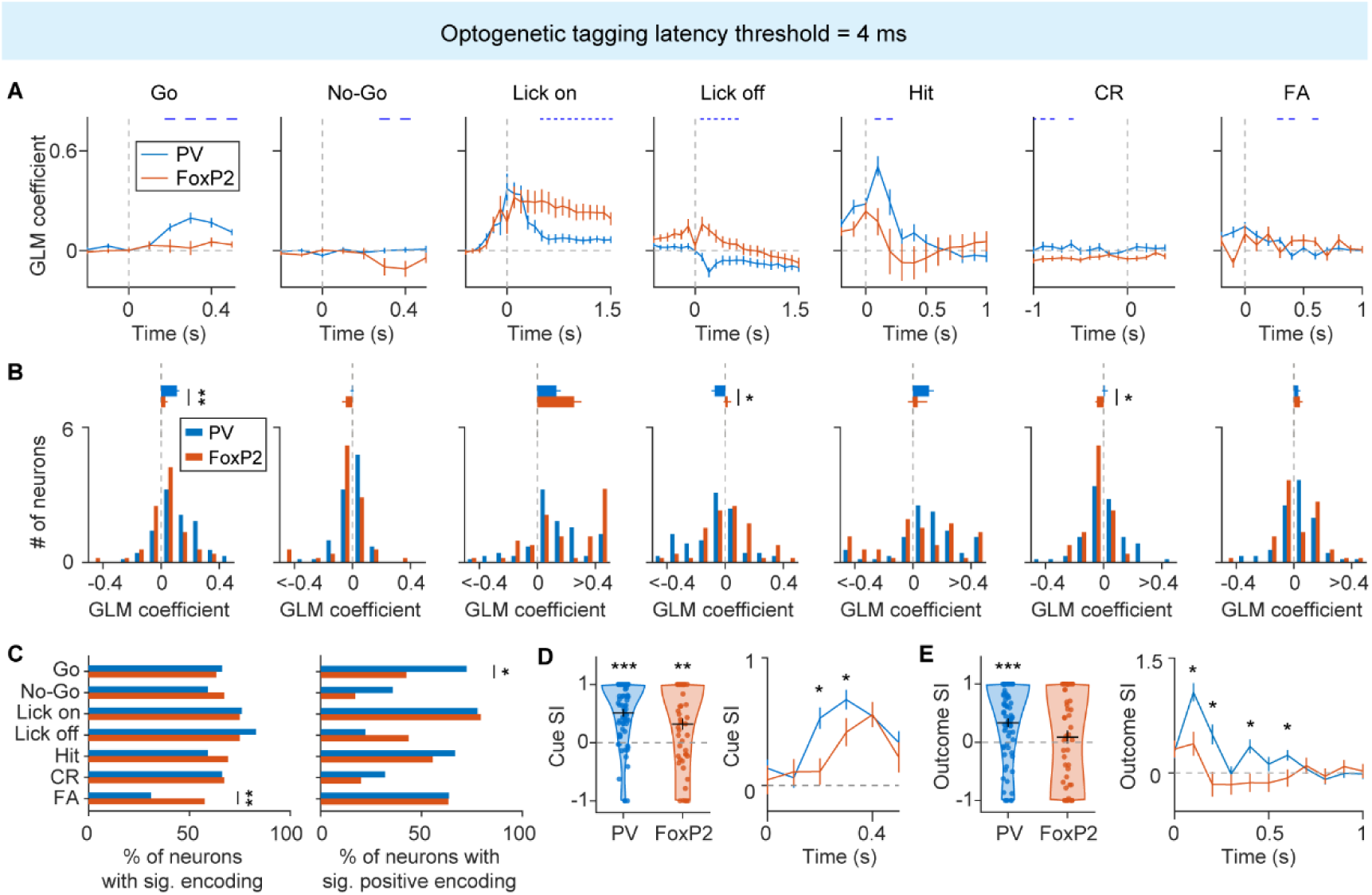
Comparison of GLM coefficients, cue SI, and outcome SI between task-modulated PV and FoxP2 GPe neurons identified using a 4-ms latency threshold. (A) Temporal profiles of GLM coefficient averaged across PV neurons (n = 71) or FoxP2 neurons (n = 52) for each regressor. Horizontal ticks indicate time points where GLM coefficients differed significantly between PV and FoxP2 neurons (p < 0.05), determined by a two-way ANOVA with mixed design followed by Tukey’s multiple comparisons test. (B) Distribution of the mean GLM coefficients averaged over the same time windows specified in (A). For Go stimulus, p = 0.0024; for lick-bout onset, p = 0.011; for lick-bout offset, p = 0.018; for Hit, p = 0.04; two-sample Kolmogorov-Smirnov test. Top, average of GLM coefficients. * p < 0.05, ** p < 0.01, Wilcoxon rank-sum test. (C) Left, proportion of PV and FoxP2 neurons showing significant encoding for each task event. Right, among neurons with significant encoding, proportion of PV and FoxP2 neurons exhibiting significantly positive (or negative) encoding for each event. * p < 0.05, ** p < 0.01, Fisher’s exact test. (D) Left, cue SI was significantly larger than 0 for both PV and FoxP2 GPe neurons (p < 0.01, Wilcoxon signed-rank test), with no significant difference between the two cell types (p = 0.26, Wilcoxon rank-sum test). Right, temporal dynamics of cue SI in PV versus FoxP2 neurons. * p < 0.05, two-way ANOVA with mixed design (F_(1, 121)_ = 4.02, p = 0.047) followed by Tukey’s multiple comparisons test. (E) Left, outcome SI of PV neurons was significantly greater than zero (p = 9.7×10^-5^, Wilcoxon signed-rank test), whereas that of FoxP2 neurons did not differ significantly from zero (p = 0.43, Wilcoxon signed-rank test). Right, temporal dynamics of outcome SI in PV versus FoxP2 neurons. * p < 0.05, two-way ANOVA with mixed design (F_(1, 121)_ = 5.94, p = 0.016) followed by Tukey’s multiple comparisons test. Data represent mean ± SEM.

**Figure 6 - figure supplement 1.**
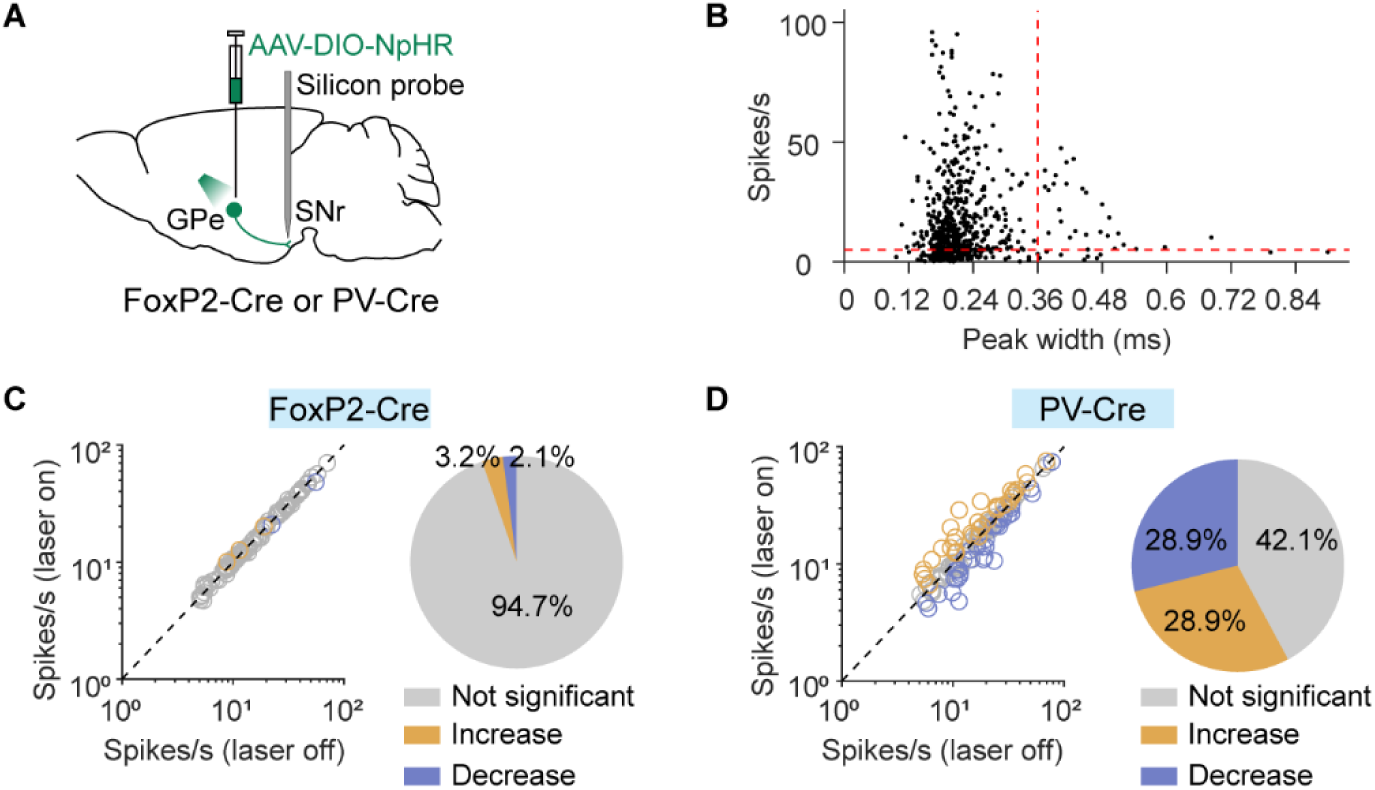
Effect of inhibiting GPe FoxP2 or PV neurons on spontaneous activity of putative GABAergic SNr neurons. (A) Schematic of SNr recording combined with GPe FoxP2 or PV neuron inactivation. (B) Putative GABAergic SNr units were identified by a spike peak width < 0.36 ms, and only neurons with firing rates > 5 Hz were included in the analyses. (C) Left, spontaneous firing rates of SNr neurons with vs. without inactivating GPe FoxP2 neurons. Right, proportion of SNr neurons showing significantly increased, decreased, or unchanged firing rates. n = 95 SNr neurons from n = 2 FoxP2-Cre mice. (D) Left, spontaneous firing rates of SNr neurons with vs. without inactivating GPe PV neurons. Right, proportion of SNr neurons showing significantly increased, decreased, or unchanged firing rates. n = 121 SNr neurons from n = 2 PV-Cre mice.

**Figure 6 - figure supplement 2.**
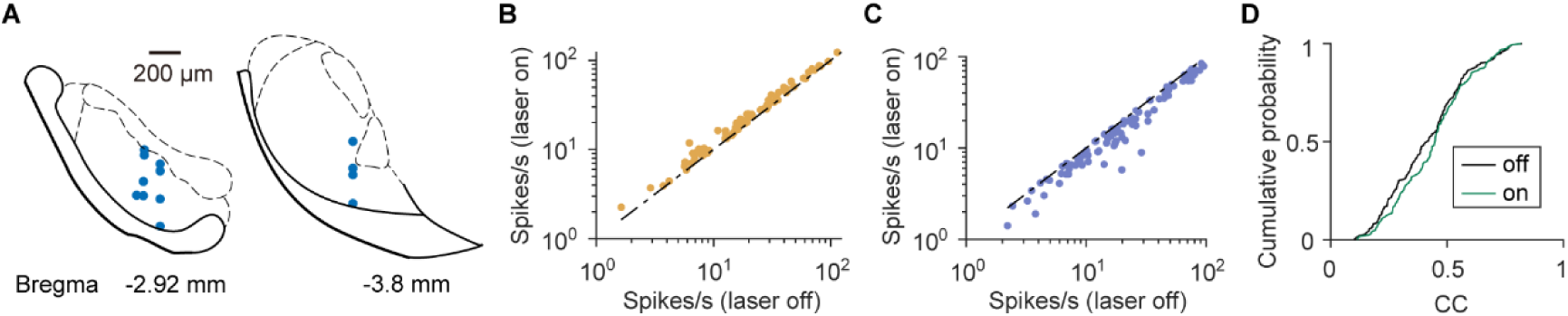
Recording sites, firing rate changes of putative GABAergic SNr neurons upon inactivation of GPe PV neurons in behaving mice, and task-related GLM performance of SNr neurons. (A) Recording locations of SNr neurons. Each dot represents the position of electrode tip in the last recording session for each mouse. Scale bars, 200 μm. (B) In a subset of putative GABAergic SNr neurons (n = 74), inactivation of GPe PV neurons significantly increased mean firing rates. The mean firing rate of each neuron was calculated from 0 – 3 s after stimulus onset, and significance was determined by comparing laser-off and laser-on trials (Go or No-Go) using a two-way ANOVA on 100-ms binned PSTHs (p < 0.01). (C) In another subset of putative GABAergic SNr neurons (n = 88), inactivation of GPe PV neurons significantly decreased mean firing rates. (D) For neurons with significant firing rate change, the correlation coefficients (CCs) between GLM-predicted and actual activity were calculated separately for laser-off and laser-on trials. The distributions of CCs under the two conditions did not differ significantly (p = 0.20, n = 162, two-sample Kolmogorov-Smirnov test).

**Figure 6 - figure supplement 3.**
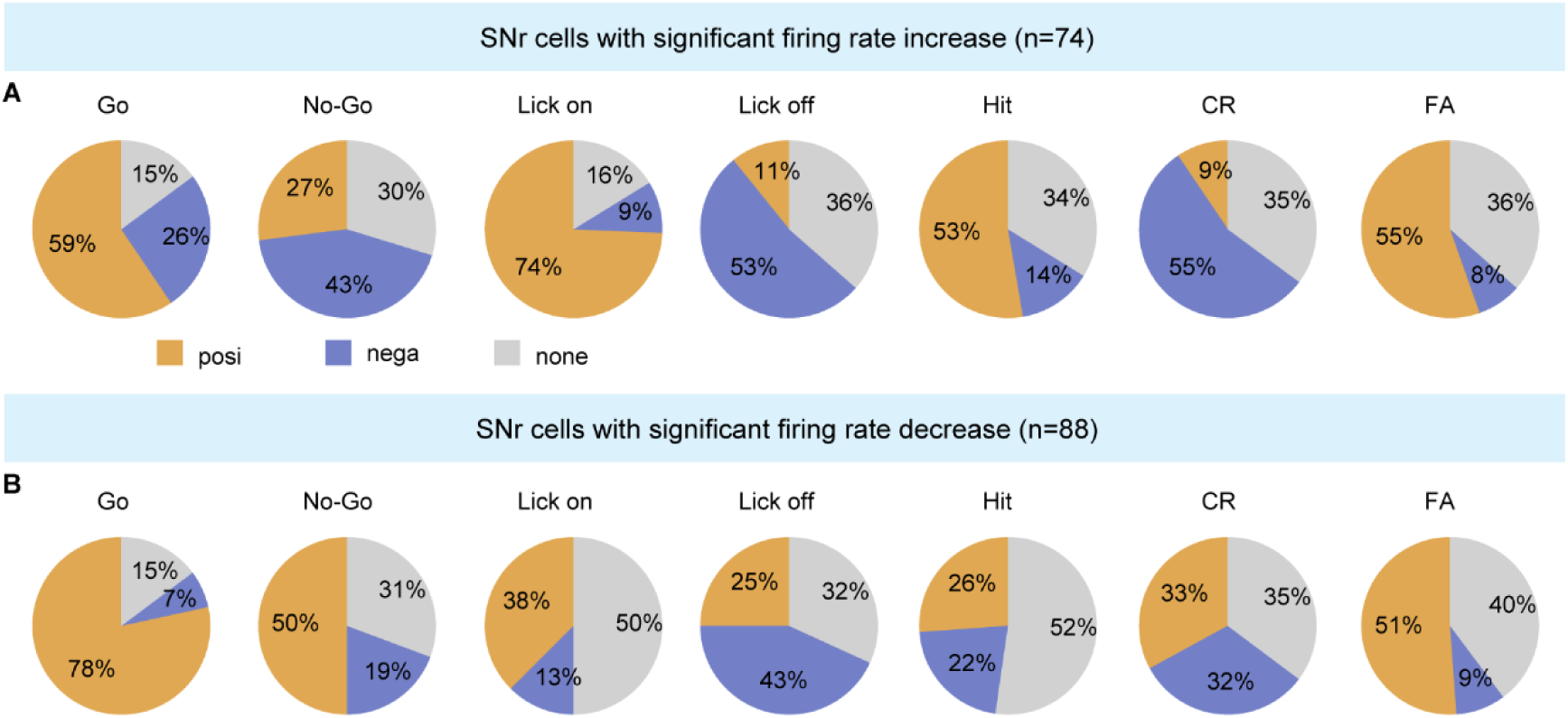
Fraction of SNr neurons with significant encoding for each event. (A) SNr neurons with significant firing rate increase upon GPe PV neuron inactivation. Each pie chart indicates the proportions of neurons exhibiting significant positive encoding, significant negative encoding, or no significant encoding for a given event. (B) Same as (A), but for SNr neurons with significant firing rate decrease upon GPe PV neuron inactivation. Among the seven task events, five (Go, No-Go, lick-bout onset, Hit, and CR) showed significant differences in the distributions of neurons with significant positive, significant negative, and non-significant encoding between firing-increased and firing-decreased SNr neurons (p < 0.01 for each event, χ^2^ test).

## Notes

https://doi.org/10.17632/z6sg8rzhxb.2

### Competing Interest Statement

The authors have declared no competing interest.

### Summary of Updates

Figure 3, Figure 5, and Figure 7 have been revised, and the Supplementary files updated. We have revised the interpretation of the selectivity index (SI) analysis, now describing SI as reflecting stimulus or outcome preference rather than neuronal selectivity. In addition, we performed in vitro slice electrophysiology to validate NpHR-mediated inhibition of GPe FoxP2 neurons and added new experiments examining the effects of inhibiting GPe PV or FoxP2 neurons on spontaneous firing in SNr neurons. The corresponding text and figures have been updated throughout the manuscript.

